# Minimal functional alignment of ventromedial prefrontal cortex intracranial EEG signals during naturalistic viewing

**DOI:** 10.1101/2021.05.10.443308

**Authors:** Tiankang Xie, Jin Hyun Cheong, Jeremy R. Manning, Amanda M. Brandt, Joshua P. Aronson, Barbara C. Jobst, Krzysztof A. Bujarski, Luke J. Chang

## Abstract

The ventromedial prefrontal cortex (vmPFC) has been thought to play an important role in processing endogenous information such as generating subjective affective meaning. Unlike sensory cortex, which processes exogenous information about the external world similarly across individuals, prior work has posited that vmPFC activity may be idiosyncratic to each individual, even when exposed to the same external stimulus. In this study, we recorded local field potentials (LFPs) from intracranial stereotactic electrodes implanted in patients with intractable epilepsy while they watched an emotionally engaging television show episode and evaluated temporal synchronization of these signals across participants in auditory cortex and vmPFC. Overall, we observed markedly lower intersubject synchronization of signals recorded from electrodes implanted in vmPFC compared to auditory cortex. A subset of patients, however, appeared to share similar vmPFC states during the more emotionally salient scenes. This work suggests that the vmPFC is involved in processing affective responses to ongoing experience in a state-like manner, but the specific states and temporal sequences are idiosyncratic to each individual, even when viewing the same television episode.

## Introduction

The ventromedial prefrontal cortex (vmPFC) is a functionally heterogeneous cortical region (de la Vega et al., 2016; Kahnt et al., 2012) that has been the subject of extensive investigation over the past few decades. It is metabolically active in the absence of any explicit task (Raichle et al., 2001) and coactivates with a distinct functional network that includes the posterior cingulate cortex (Buckner and DiNicola, 2019; Fox and Raichle, 2007). The vmPFC plays a central role in learning, memory, and decision-making by facilitating the integration of multimodal value signals (Bartra et al., 2013; Hare et al., 2008; Knutson et al., 2001; Padoa-Schioppa and Assad, 2006; Rangel et al., 2008; Rich and Wallis, 2016; Suzuki et al., 2017), representing latent contextual information (Constantinescu et al., 2016; Niv, 2019; Schuck et al., 2016), and remembering the past and projecting into the future (Buckner and Carroll, 2007). The vmPFC also processes affective and social information by generating (Chikazoe et al., 2014; Damasio et al., 2000; Eisenbarth et al., 2016; Lindquist et al., 2012) and regulating affective states (Etkin et al., 2015; Ochsner et al., 2002; Phelps et al., 2004; Wager et al., 2004), and by supporting internally generated self-referential thought (Andrews-Hanna et al., 2014; Kelley et al., 2002; Mason et al., 2007; Mitchell et al., 2002). Because the vmPFC is anatomically connected to many systems involved in processing affective and conceptual information including the brainstem, insula, medial temporal lobe, and prefrontal cortex, it has been hypothesized to be the hub in generating affective meaning (Ashar et al., 2017; Chang et al., 2021; Damasio, 2006; Roy et al., 2012). However, generating affective meaning is a highly idiosyncratic process that synthesizes endogenous signals that are unlikely to be common across participants, even when they are exposed to the same exogenous stimuli or experiences. This means that the same person might generate different affective meaning from others in response to the same eliciting event based on fluctuations in these endogenous signals.

Unlike unimodal cortex that processes exogenous information about the external world, the vmPFC is at the pinnacle of transmodal association cortex that processes endogenous information pertaining to our past experiences, current homeostatic states, and future goals (Margulies et al., 2016; Margulies and Smallwood, 2017; Mesulam, 1998; Paquola et al., 2019). Consequently, responses in this region are highly variable across individuals, exhibiting little evidence of intersubject spatiotemporal synchronization across a variety of experimental contexts such as listening to stories (Lerner et al., 2011), watching movies (Chang et al., 2021; Chen et al., 2017; Hasson et al., 2004), multivariate decoding (Bhandari et al., 2018), or functional connectivity patterns in resting state fMRI (Gordon et al., 2016; Mueller et al., 2013). For example, in a recent fMRI study, in which participants watched a pilot episode from a character-driven television drama, we observed minimal amounts of intersubject synchronization in the vmPFC compared to unimodal sensory cortex (Chang et al., 2021). Instead, participants appeared to dynamically switch between a few discrete states while watching the show. Though the vmPFC states were spatially consistent across participants, they were experienced at different times for each participant, with the largest temporal synchronization of states occurring at the most emotionally salient moments in the episode. Using these spatial patterns as indicators of an endogenous state revealed consistent activations distributed throughout the brain in distinct affective networks despite not being consistently time-locked to the external stimuli. While this study demonstrated a clear dissociation between how exogenous and endogenous information are processed in the brain, there are a number of open questions that remain unanswered. First, though the primary results were replicated across two independent fMRI datasets, it is unclear how much the effects might simply reflect artifacts of the BOLD fMRI signal itself. For example, the vmPFC is notoriously difficult to image due to its close proximity to the orbital sinus, which creates susceptibility artifacts in the magnetic field resulting in large signal dropout and spatial distortion (Glover and Law, 2001; Weiskopf et al., 2007). Second, the BOLD signal reflects a downstream process of neural firing, and heterogeneity in individual vascular systems may lead to individual variations in hemodynamic responses (Birn et al., 2001; Handwerker et al., 2012; Lindquist et al., 2009).

To address these limitations, we sought to more directly assess temporal synchronization of vmPFC activity using intracranial electrophysiological recordings of local field potentials (LFPs). Unlike fMRI, intracranial recordings do not suffer from susceptibility artifacts, and can measure signals with high temporal resolution from very specific spatial locations. Prior intracranial work investigating the vmPFC has relied on more traditional task-based paradigms to characterize its role in computing value (Hill et al., 2016; Li et al., 2016; Lopez-Persem et al., 2020; Saez et al., 2018). Stimulation paradigms have implicated the involvement of the vmPFC in the subjective experience of affect, olfaction, and gustation (Fox et al., 2018; Yih et al., 2019). Naturalistic designs, such as passive movie watching, can provide rich contextual information, which makes them well suited for studying social, cognitive, and affective processes (Hasson et al., 2020; Jolly and Chang, 2019; Sonkusare et al., 2019), but have been rarely used in intracranial EEG research (Honey et al., 2012; Jafarpour et al., 2019; Mukamel et al., 2005). The majority of work using naturalistic designs has instead focused on characterizing sensory processes in auditory cortex and found that power in high frequency bands, such as gamma (***γ***), positively correlate with the envelope of the auditory stimulus (Honey et al., 2012) and also with fluctuations in the BOLD response in auditory cortex of other participants viewing the same stimulus while undergoing fMRI (Mukamel et al., 2005). Power in low frequency bands (e.g., α), in contrast, exhibits an inverse relationship and negatively correlates with auditory signals and fluctuations in BOLD.

In the present study, we recorded intracranial stereo electroencephalogram (sEEG) data from 6 patients undergoing surgical intervention for intractable epilepsy while they viewed a 45-minute television episode (*Friday Night Lights)*. We were primarily interested in evaluating the temporal consistency of responses in the vmPFC across patients compared to the auditory cortex. Intersubject correlation analysis (ISC) (Hasson et al., 2004; Nastase et al., 2019) has emerged as the predominant method to evaluate the functional alignment of signals across participants. High levels of ISC indicate that participants are processing information pertaining to the stimuli similarly, while low ISC indicates high intersubject heterogeneity. However, there are several difficulties applying this technique to sEEG data. First, both the number and location of implanted sEEG electrodes are determined by clinical needs rather than research interests, which results in only a small fraction of the cerebral cortex being covered by any given patient’s electrodes, along with little overlap in implantation locations across patients. Unlike electrocorticography (ecog) recordings in which a high density grid of electrodes is placed over the same cortical surface, across-subject comparisons are particularly difficult with sEEG as there can be large inter-subject differences in the number and location of electrodes (Chang, 2015; Owen et al., 2020; Parvizi and Kastner, 2018). Second, electrodes implanted in the same participant and recorded from the same stereotactic strip are not entirely independent, and are likely to be reflecting similar signals. Failure to account for these within-subject clustering effects and electrode distances can bias the ISC metric. To address these limitations, we assess the impact of within subject similarity and spatial distance on ISC in the vmPFC and auditory cortex and evaluate the ability of functional alignment (Chen et al., 2015a; Haxby et al., 2020) to overcome these inherent challenges to working with sEEG data.

## Results

### Inter-electrode temporal synchrony

To estimate and quantify the influence of spatial distance, within-subject, and within-strip clustering on inter-electrode similarity, we fit a linear distance regression model separately to vmPFC and auditory electrodes broadband power activities. Specifically, we predicted the pairwise inter-electrode broadband power temporal similarity matrix from a linear combination of: (a) the inter-electrode spatial distance, (b) dummy variables indicating which pairs of electrodes belonged to each of the 6 participants, and (c) dummy variables indicating which pairs of electrodes belonged to the same electrode strip (Fig. 2A). This analysis allows us to quantitatively model the independent effects of spatial distance and within-subject effects on inter-electrode temporal synchronization. In vmPFC electrodes, we found that spatial distance (F(1) = 5.28, p = 0.0218, r^2^ = 0.0001), within-subject effects (F(7) = 3497.7, p < 0.001, r^2^ = 0.535) and within-strip effects (F(16) = 85.67, p < 0.001, r^2^ = 0.030) all independently explained a substantial portion of inter-electrode similarity variance beyond inter-electrode clustering within the same stereotactic strip. Similarly, for auditory electrodes we found that spatial distance (F(1) = 7.02, p = 0.008, r^2^ = 0.0004), within-subject effects (F(5) = 843.77, p < 0.001, r^2^ = 0.224), and within-strip effects (F(9) = 97.94, p < 0.001, r^2^ = 0.047) all significantly accounted for temporal synchrony variance (Fig. 2D).

These linear distance regression models allowed us to quantitatively remove this nuisance variance from the pairwise temporal similarity matrix. We computed ISC on the residuals after removing the effects of spatial distance and within-subject effects and observed higher levels of residual inter-electrode temporal synchrony in auditory cortex (r=0.014, bootstrap p=0.013) compared with vmPFC (r=0.003, bootstrap p=0.381) using a sign permutation test (p<0.001; Fig. 2C). We also examined these effects across multiple frequency bands and consistently found low ISC values in vmPFC across frequency bands (Fig. S2).

Taken together, these results indicate that within both vmPFC and auditory cortex, broadband temporal dynamics were more similar across electrodes within the same participant and within the same stereotactic strips than across participants, and that nearby electrodes exhibited more similar broadband temporal dynamics than distant electrodes. After conditioning on all of these effects, we found that electrodes within the auditory cortex exhibited significantly higher levels of broadband temporal synchronization across subjects than electrodes within vmPFC.

### Improving inter-electrode alignment with the Shared Response Model

Our distance regression approach allowed us to estimate the impact of spatial distance and the subject-specific correlation pattern for each participant (Fig. 1, diagonal blocks). However, these regressions are based on the assumptions that: (a) millimeter-scale discrepancies in spatial location correspond to linear changes in temporal synchrony within each region, and (b) each electrode is statistically independent of the other electrodes. Although these simplifying assumptions enabled us to gain the new insights summarized above, we also know from prior work that neither of these assumptions are likely to be (strictly) true. To address these issues, we used a completely different analytic approach and evaluated the efficacy of aligning the electrodes across participants within each region to a common latent space using the Shared Response Model (SRM) (Chen et al., 2015a).

**Figure 1.**
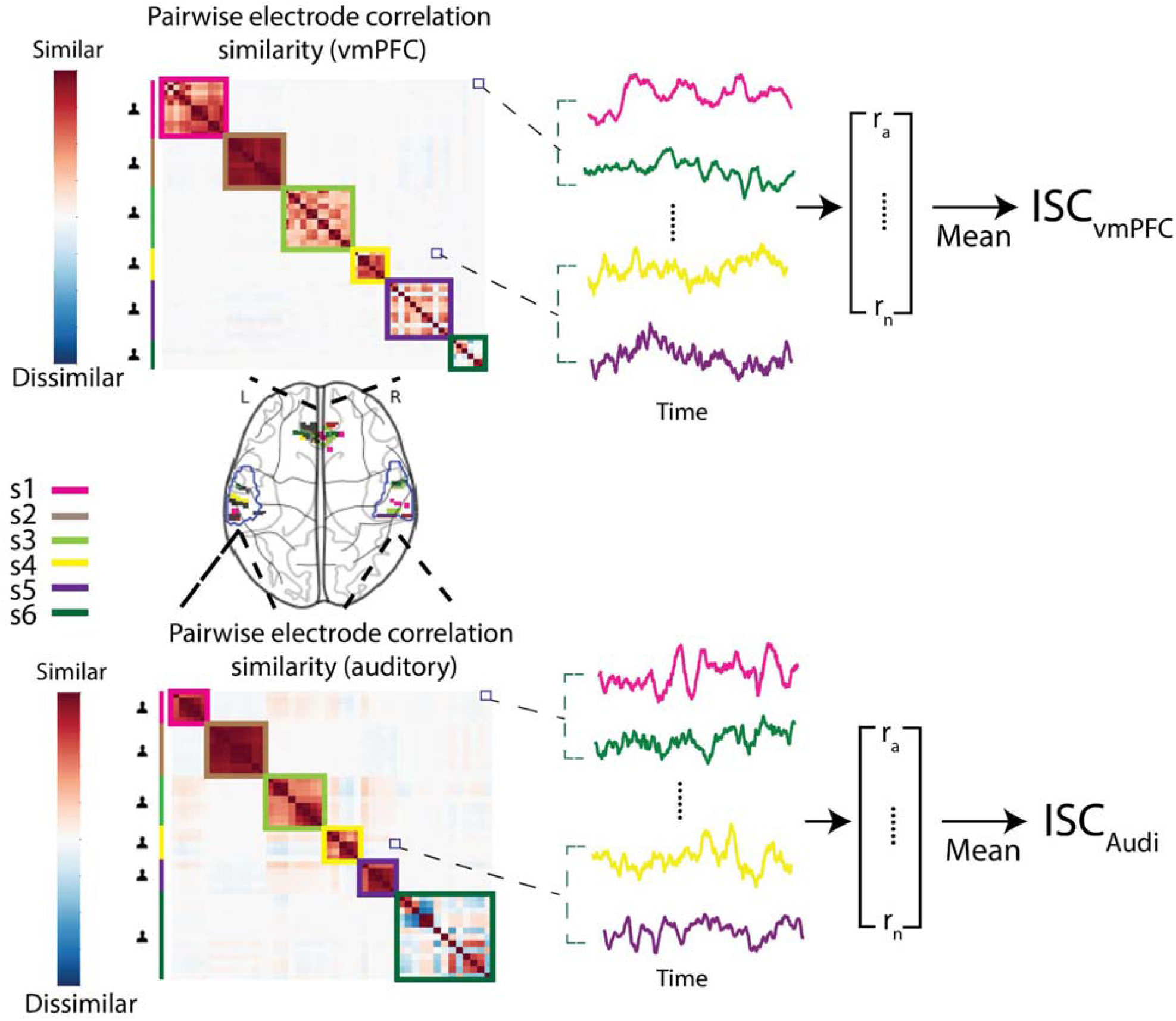
Electrode locations and temporal similarity. Here we illustrate the electrode locations within the vmPFC and bilateral auditory regions of interest. Electrode anatomical locations vary across subjects. We compute the pairwise correlations between electrodes. The similarity of electrode pairs recorded from within the same participants is considerably higher than across participants. Each subject is indicated by a unique color. Intersubject Correlation (ISC) is the mean of the lower triangle of this pairwise correlation matrix after performing a Fisher r to z transformation.

**Fig. 2.**
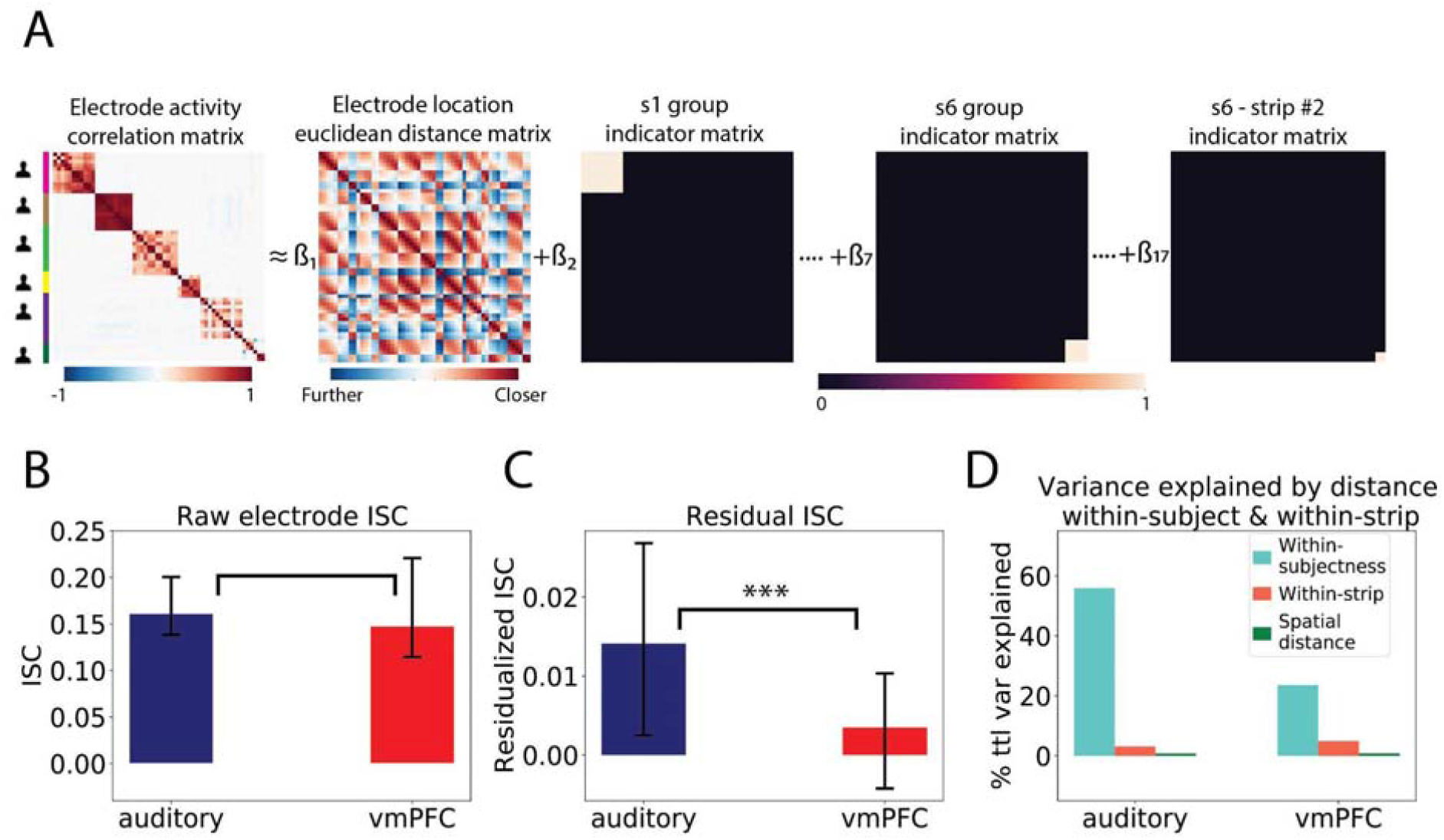
Temporal Synchrony Distance Regression. A. We used a fixed-effects regression to separate out the effect of spatial locations and within-subject clustering on pairwise electrode temporal distance. B. We observed high temporal synchronization of broadband power in both vmPFC and auditory electrodes. The auditory broadband ISC is not statistically different from the vmPFC broadband ISC (sign permutation test, p = 0.21). C. After removing within-subject clustering and spatial distance effects, we see a marked decrease in temporal ISC in vmPFC, but not auditory cortex. Auditory broadband ISC is significantly higher than vmPFC (sign permutation test, p < 0.001) D. Overall percentage of variance explained by spatial distance and within-subject clustering in auditory/vmpfc regression model (i.e., r^2^) normalized by the total variance explained by the full model. Although both explain a significant amount of variance, the within clustering effect explains the majority of variance in the pairwise correlation values. Error bars indicate 95% confidence interval for the ISC values (See details in Method section).

SRM was originally developed to perform functional alignment (Haxby et al., 2020, 2011) on fMRI data by remapping voxels into a common latent space based on shared responses to time-synchronized stimuli (Chen et al., 2015a; Vodrahalli et al., 2018). This mapping is learned using a data-driven unsupervised latent factor algorithm and allows each subject to have a different number of electrode recordings. Formally, for each subject ***i***’s number-of-electrodes by number-of-timepoints data matrix,***X***_***i***_, SRM finds an individual basis ***W***_***i***_, and a latent time series matrix ***S***, subject to the constraint that ***S*** is common across all participants. This allows ***X***_***i***_ to be approximated by ***X***_***i***_ = ***W***_***i***_***S + E***_***i***_, where ***W***_***i***_ and ***E***_***i***_ are subject-specific and where ***S*** is the latent embedding shared across all ***m*** participants (Fig. 3). Here, we use SRM to align electrodes separately within the auditory or vmPFC regions after performing minimal preprocessing (i.e., bandpass filtering, bad channel removal, and re-referencing), based on putative shared temporal dynamics across participants. The output of this procedure is a separate component-by-time matrix for each participant, in which each component is now aligned across participants and can be used in place of voltage in any other type of analysis.

**Fig. 3.**
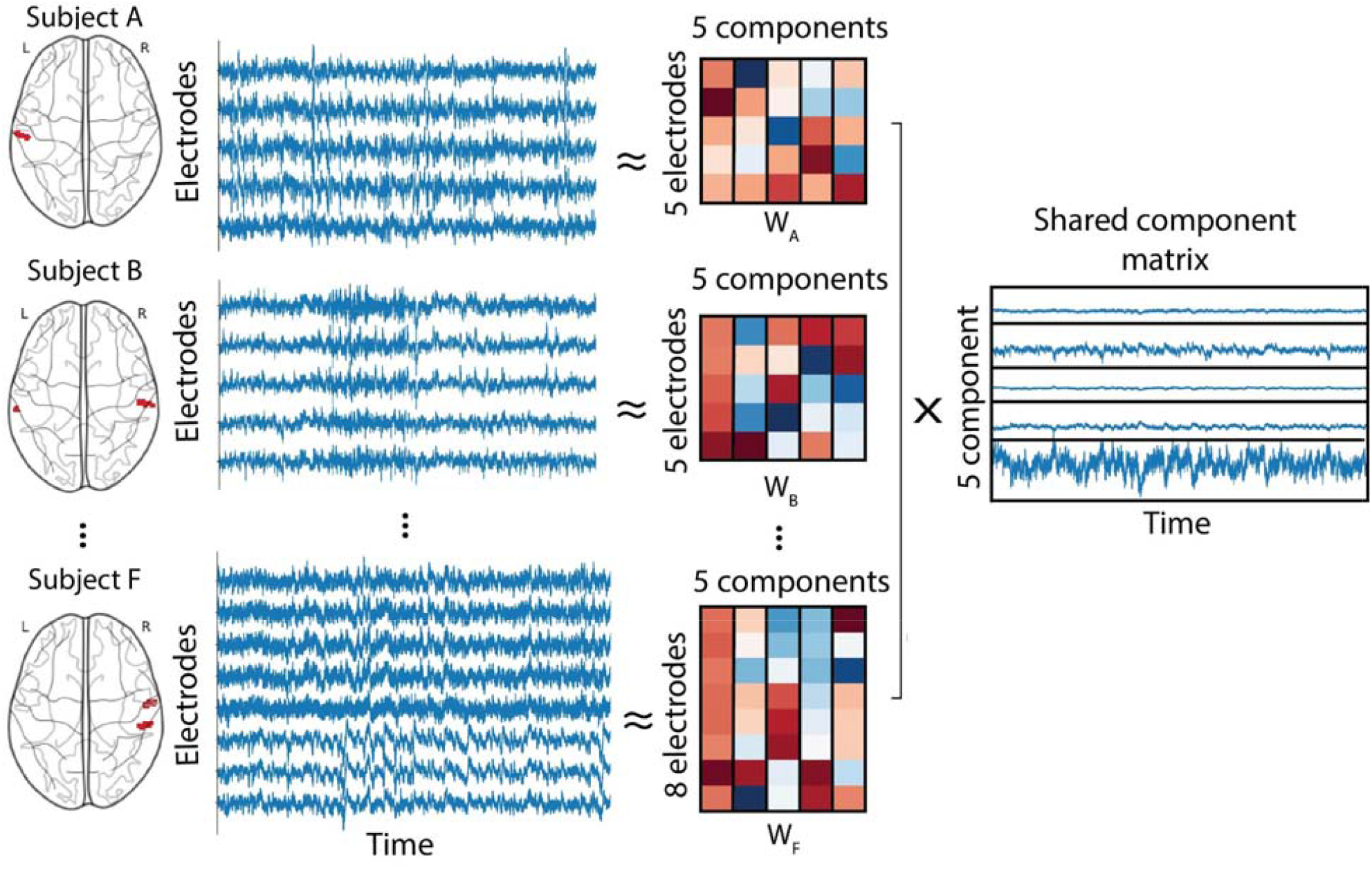
Shared Response Model. Graphical depiction of Shared Response Model applied to auditory electrodes. We applied SRM to electrodes in 6 subjects and decomposed electrode voltage in each subject into subject-specific basis functions that project into a shared component matrix common across all subjects.

We first attempted to validate how well this technique performs in functionally aligning the sEEG electrodes. We hypothesized that functional alignment should aid in extracting functional signals that are common across participants filtering out idiosyncratic signals. Specifically, we anticipated that aligning electrodes within the auditory cortex should result in a signal that better corresponds to properties of the auditory stimulus while participants watched the show compared to simply computing the mean across electrodes. For example, prior work has shown that lower frequency power tends to negatively correlate with the audio envelope of auditory stimuli, whereas higher frequency power tends to positively correlate with the auditory signal (Honey et al., 2012; Mukamel et al., 2005; Nir et al., 2007). To test this hypothesis, we estimated a 5-dimensional SRM using the band-pass filtered voltage activities of 44 auditory electrodes from 6 subjects. We selected the SRM component with the highest ISC across subjects (ISC = 0.04, p<0.001) and computed the time-varying power in the low ***δ*** (i.e., 1-2 Hz) and high ***γ*** (i.e., 70-150 Hz) power bands. We then correlated these signals with the raw audio envelope to assess how well this approach was able to recover the true signal we expected to be present in the data (Fig. 4A). Consistent with prior work (Honey et al., 2012; Mukamel et al., 2005), we found that power in the ***δ*** band negatively correlated with the audio envelope (r=-0.22, p<0.001), while the broadband ***γ*** power of the SRM component positively correlated with the raw audio envelope (r=0.06, p<0.001). Importantly, we also found that the SRM component exhibited a tighter coupling of the audio envelope in both ***δ*** and high ***γ*** frequency bands compared to simply averaging power from electrodes across participants (low delta: p < 0.001; broadband gamma: p = 0.02). This analysis provides a proof-of-concept demonstration confirming that the SRM can extract meaningful functional signals from a well-characterized region of cortex, and outperforms across-participant electrode averaging.

**Fig. 4.**
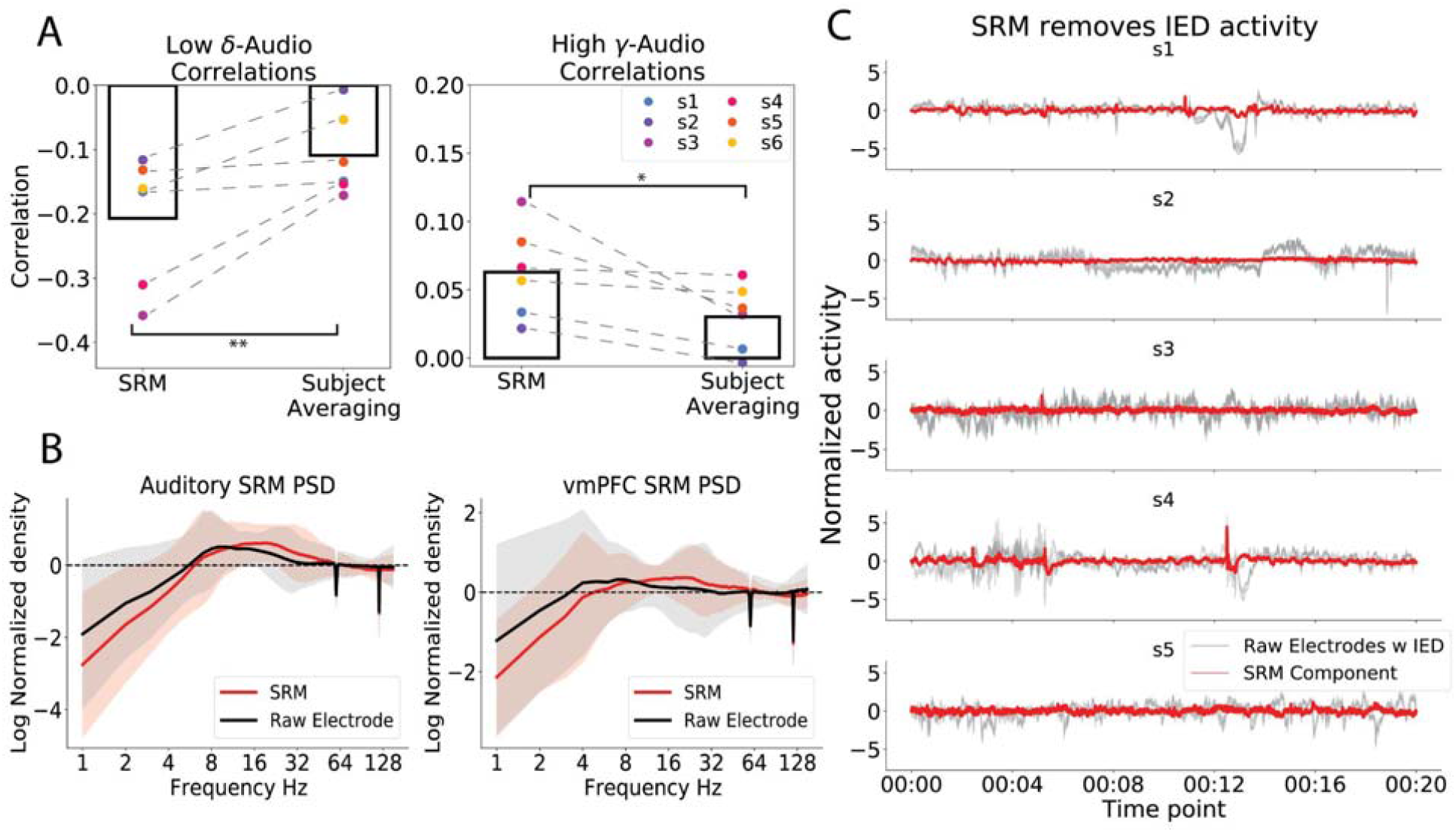
Validating SRM on auditory cortex. A: SRM component from auditory cortex electrodes better captures the audio-specific stimuli than the subject-averaging method. Left: correlation between SRM low ***δ*** power (1-2 Hz) and audio envelope; Right: correlation between SRM high ***γ*** power (70-150 Hz) and audio envelope (*: 0.01<=p<0.05; **: 0.001<p<=0.01). B: The log-log plot for Power Spectrum Density of auditory and vmPFC max SRMs. The 1/f trend is removed by fitting a first-order linear regression and obtaining the residuals. We plotted the subject-averaged and 1/f detrended PSDs for both the max SRM component (red lines) and the original electrode channels (black lines). Shaded areas indicate 95% confidence intervals. We see that the SRM does not appreciably distort the original electrode PSD. C. To estimate the extent to which SRM can reduce IED and Ictal (seizure) activities, we apply SRM to 20s of selected time periods in amygdala electrodes in 5 subjects where a mixture of IEDs and Ictal activity patterns is clearly visible by eye. We apply SRM to data with IED and Ictal activities and we found that SRM removed the subject-specific IED and Ictal activities.

We next explored the impact of the SRM transformation on the power spectral density (PSD). EEG LFPs tend to reflect higher power in lower frequency components (the well characterized 1/f noise) and this may have impacted the functional alignment procedure. We computed the mean-normalized power spectrum density (PSD) of the maximum SRM component and compared this to the PSD of the average electrode bandpass filtered voltages. Overall, we found that the SRM component did not appear to substantially alter the PSD in either auditory cortex or vmPFC (Fig. 4B; see S8, for individual participants).

Another potential benefit of applying SRM to LFPs relates to artifact removal. Because SRM attempts to temporally align signals that are shared across participants, it likely implicitly removes artifacts that are not time-locked across participants. For example, removing ictal activity and interictal epileptiform discharges (IEDs) is well-known in the intracranial literature to present a substantial computational challenge (Keller et al., 2010; Thomas et al., 2018). Several factors, including a lack of precise and established definitions of epileptic spikes and wide spectrum of morphologies of epileptiform abnormalities both within the same patient and across patients, contribute to the difficulties in identifying and removal of IEDs (Dümpelmann and Elger, 1999; El-Gohary et al., 2008; Yadav et al., 2011). However, because IED morphologies are unique to each participant and occur at different times, we hypothesized that SRM should be effective in removing these idiosyncratic signals. To test this hypothesis, we estimated a new SRM using the non-preprocessed raw voltage data and found that SRM was able to successfully remove IEDs and Ictal activities (Fig. 4C). In addition, SRM was able to reduce the impact of 60Hz power line noise by an average of 51% (Fig. S3; see supplementary materials for details).

### vmPFC activity does not align across subjects

After validating the SRM method on the auditory cortex, we next were interested in evaluating shared signals in the vmPFC after aligning electrodes with SRM. We trained an SRM with 6 components on vmPFC electrodes and transformed each participant’s observed data into the shared latent space. We selected the SRM component with the highest ISC across subjects (ISC=0.01, p=0.004). We also computed the ISC across broadband power and several narrow band frequencies (i.e., ***δ, θ, α, β***, and low and high ***γ***). We did not observe a significant effect of intersubject synchronization across any frequency band in the vmPFC using a bootstrapping procedure (Fig. 5; ***δ***: r=0.01, p = 0.08 **θ**: r=0.01, p = 0.95 ***α***: r=-0.00, p = 0.93 ***β***: r=0.00, p = 0.24 low ***γ***: r=0.00, p = 0.12 high : r=0.00, p=0.27, broadband: r=0.01, p = 0.49), unlike auditory cortex (Fig. 5; ***δ***: r=0.11; p = 0.002; **θ**: r=0.02, p = 0.16; ***α***: r=0.06; p < 0.001 ***β***: r=0.024, p = 0.07; low : r=0.01, p = 0.23; high ***γ***: r=0.01, p=0.1; broadband: r=0.03, p = 0.02) Moreover, we found that the ISC in the SRM-aligned vmPFC was significantly reduced compared to auditory cortex across multiple bands using a permutation test (***δ***: effect size diff = 0.09, p < 0.001, ***α***: effect size diff = 0.06, p < 0.001, ***β***: effect size diff = 0.02, p = 0.002, broadband: effect size diff = 0.02, p = 0.003). To ensure that these results were completely unbiased from training and testing the SRM model on the same data, we also performed a split-half cross-validation procedure, in which we trained the model on the first or second half of the data and tested it on the other half. Overall, we observed consistent results, in which ISC was not significantly different from zero, even when the SRM model was trained on independent data (Fig. S4).

**Fig. 5.**
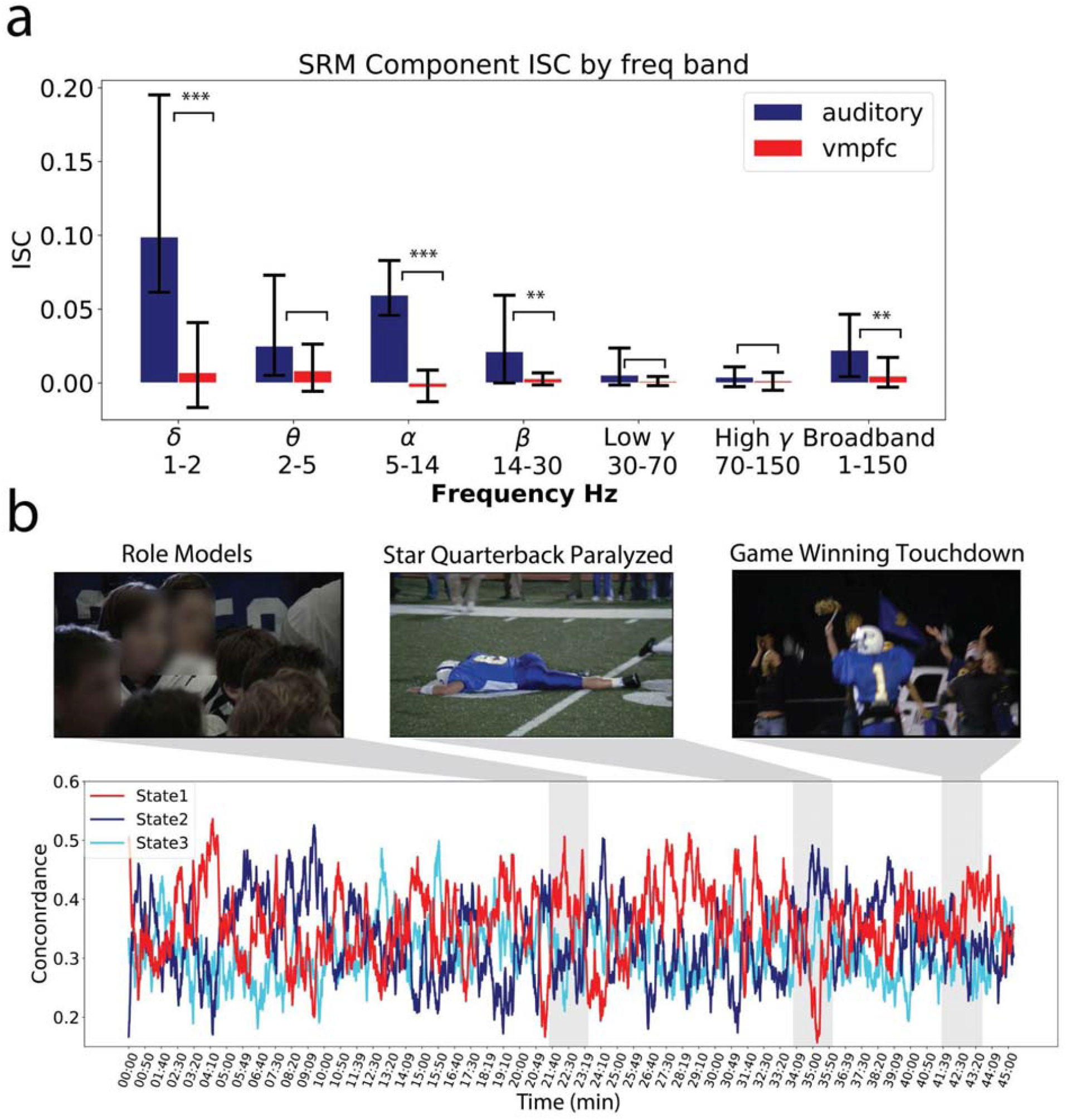
Shared Response Model vmPFC Alignment. A) Inter-subject correlation value for auditory (blue) and vmPFC (red) electrodes. After identifying the SRM component with the highest ISC across participants, we extracted power from different frequency bands and computed ISC within each band including broadband power. Error bars indicate 95% confidence interval for the ISC values (See details in Method section), * indicates p<0.05 using subject-wise bootstrapping. B) Line plots illustrate each HMM state concordance across participants at each time moment during the episode and have been smoothed with an exponential smoothing function with alpha = 0.001 for visualization. Shaded regions indicate scenes associated with intense affective experiences reported in (Chang et al., 2021). State 1 appears corresponds to positive affective scenes while State 2 corresponds to negative affective scenes. Screenshots from Friday Night Lights are copyright of NBCUniversal, LLC.

### Temporal alignment of discrete SRM states across participants

The above ISC analyses provide converging evidence with the distance regression analyses indicating minimal evidence of consistent across-participant temporal synchronization in the vmPFC across the 45min episode consistent with our hypotheses. However, there are many other possible explanations for this finding including that our SRM alignment approach may have inadvertently removed much of the signal of interest and we are simply modeling the residual noise. One proposed approach to address this is to have participants view the same stimulus twice and compute the intra-viewing synchronization coefficient within each participant (Nastase et al., 2019). However, this approach is problematic for studying processes beyond sensory perception such as memory or generating affective meaning as the second viewing will necessarily be impacted by the first viewing, which will decrease within-subject ISC values. Instead, we used an entirely different approach to evaluate if there might be particular time points during which participants briefly experienced the same psychological state. This would demonstrate that: (a) we have retained true signal, and also (b) provide evidence of inter-subject idiosyncratic processing. For example, in a previous fMRI study, we found evidence that participants were more likely to occupy the same psychological state during scenes that elicited more intense affective experiences (Chang et al., 2021). These states were identified directly from brain activity using Hidden Markov Models (HMMs), and were validated on self-reported time-varying feelings and facial expressions.

Using this same time-varying latent state approach, we fit an HMM to the broadband power of the 6 SRM components separately for each individual participant. We made the following implicit assumptions: (1) the latent state transitions follow a first order Markovian Process, (2) the SRM components were modeled using an orthogonal multivariate Gaussian Distribution, and (3) all participants experienced the same number of latent states. We used the Viterbi algorithm (Forney, 1973) to obtain the most likely sequence of latent states for each participant. Hidden states derived from each HMM were then aligned across participants by maximizing the cross-subject state similarity using the Hungarian Algorithm (Kuhn, 2005). We estimated the number of HMM states by computing the Bayesian Information Criterion (BIC) across a range of states (***k* = [2**,**15]**) and found that k=3 states exhibited the greatest improvement in model fit (Fig. S7). Finally, we computed the across-patient state concordance at each moment in time by calculating the proportion of participants occupying the same state within each time interval (Fig 5B). Higher concordance values indicate that more participants were sharing a common psychological state.

During several scenes that we identified in our prior work as having particularly high emotional salience and narrative importance (Chang et al., 2021), we found that the patients tended to converge on the same HMM states. These scenes included: (a) a positive sentimental scene when the football players provide mentorship to younger kids (∼22 min, State 1), (b) a negative scene when the star quarterback is severely injured during a play and undergoes emergent spinal surgery (∼35 min, State 2), and (c) a positive scene when the nervous and inexperienced backup quarterback throws a game winning pass (∼42 min, State 1). Though only 50% of the patients converged on the same states during these scenes, it is notable that State 1 concordance increased during positively valenced scenes, while State 2 concordance increased during negative arousing scenes similar to what we found using fMRI (Chang et al., 2021). These results are consistent with the hypothesis that the vmPFC is involved in generating affective meaning (Ashar et al., 2017; Chang et al., 2021; Chikazoe et al., 2014; Roy et al., 2012) and importantly demonstrate that the minimal synchronization observed in the ISC analyses cannot be explained by the absence of any meaningful signal in the vmPFC LFPs.

## Discussion

The majority of cognitive neuroscience research has focused on mapping stimulus-driven neural activity patterns that are common across participants. This approach has been highly successful in developing a deep understanding of how the brain processes exogenous information about the external world. However, this approach may have limited utility when characterizing regions that process endogenous information that may be idiosyncratic to each individual participant such as transmodal cortex (Mesulam, 1998). The vmPFC has been theorized to be intimately involved in generating affective meaning by integrating stimulus-driven information with an individual’s unique history of past experiences, internal homeostatic states, and future goals (Ashar et al., 2017; Chang et al., 2021; Roy et al., 2012). Consequently, activity in this region has been highly variable across individuals in a variety of tasks (Bhandari et al., 2018; Hasson et al., 2004; Mueller et al., 2013). Consistent with these observations, we have previously found using fMRI that unlike unimodal sensory cortex, the vmPFC does not appear to synchronize across participants when passively viewing an emotionally engaging television drama *except* during particularly salient emotional scenes (Chang et al., 2021). The vmPFC, however, is notoriously difficult to image using BOLD fMRI due to signal drop out and geometric distortions arising from susceptibility artifacts. Therefore, in this paper, we sought to examine inter-subject synchronization of vmPFC activity based on LFPs recorded using sEEG in patients undergoing surgical intervention for intractable epilepsy when watching an emotionally arousing naturalistic stimulus. Overall, our results are highly consistent with the previously reported fMRI findings (Chang et al., 2021). We find strong evidence of common signals being processed in the unimodal auditory cortex, but minimal evidence of cross-participant temporal synchronization in the vmPFC across any specific frequency band. However, this region does appear to reflect the idiosyncratic ways we assign affective meaning to incoming stimuli as approximately 50% of our sample appeared to be sharing a similar valenced interpretation of the most emotionally salient scenes based on our individual-HMM analysis.

A methodological challenge to evaluating shared signals across participants using sEEG is that electrodes are not located in the same spatial locations across participants. Electrode locations are determined based on clinical needs for monitoring epileptiform activity and surgical planning (Chang, 2015; Parvizi and Kastner, 2018). For example, even though we collected data from 14 patients, only 6 had electrodes placed within both of our target regions of interest (i.e., vmPFC & auditory cortex). A major contribution of this work is the application of alignment procedures that allow signals within a region originating from different anatomical locations to be compared across participants. We used two different analytic approaches that are novel to analyzing sEEG data. First, we performed linear distance regression to estimate and remove variance in the pairwise electrode temporal similarity matrix resulting from variations in the spatial distance of electrode placement, intra-subject covariance from electrodes located on the same stereotactic strip. Overall, we found that over 95% of the variance of the electrode temporal similarity matrix could be explained by these three different types of signals. Removing these signals revealed that processes in the auditory cortex, but not vmPFC appeared to be shared across participants. Second, we performed functional alignment using SRM. This approach attempts to align latent signals present in the electrodes based on commonality across participants and importantly can accommodate different numbers of electrodes for each participant. We demonstrate that this approach is better able to recover auditory signals in the auditory cortex compared to simply averaging electrodes. In addition, SRM can also be effective in removing many sources of noise that are idiosyncratic to each individual participant such as IEDs and Ictal activities and even power line noise to some extent. This was confirmed by selectively adding different types of noise to each participant and evaluating how much of the noise was removed by the SRM procedure. Similar to the distance regression approach, we only observed common signals in the auditory and not vmPFC cortex across participants after performing this alignment procedure. These results indicate that SRM provides a promising technique for both denoising signals, but also in functionally aligning electrodes and may complement alternative techniques such as SuperEEG (Owen et al., 2020).

Our results suggest that processes in transmodal cortex such as the vmPFC appear to be idiosyncratic to each individual and do not directly map onto processing exogenous information from the eliciting naturalistic stimulus. These results are in line with our previous fMRI findings in which two independent samples of participants watched the same video (Chang et al., 2021). Importantly, this work confirms that the fMRI results cannot be fully attributed to signal dropout and geometric distortions in the BOLD signal arising from susceptibility artifacts in the magnetic field where tissue borders air in the orbital sinus. How then should this null result be interpreted? On the one hand, variations in LFPs in the vmPFC across participants could arise from participants engaging in stimulus-independent thought such as mind wandering (Christoff et al., 2016; Mason et al., 2007). Alternatively, participants could be making stimulus-dependent evaluations about the affective meaning of the stimulus with respect to their idiosyncratic goals, experiences, and homeostatic states (Ashar et al., 2017; Chang et al., 2021; Roy et al., 2012). One of the most simple appraisals an individual can make is whether they like or dislike a stimulus or event. These subjective value judgments require integrating multiple attributes of a stimulus with respect to salient goals (Padoa-Schioppa and Assad, 2006; Rangel et al., 2008). For example, when appraising the value of food, the vmPFC appears to integrate multiple attributes such as taste, cost, and caloric content (Suzuki et al., 2017) with broader goals such as eating healthy (Reber et al., 2017) and internal homeostatic drives (Robinson and Berridge, 2013) and neuronal firing in this region appears to reflect the deliberation when making these decisions (Rich and Wallis, 2016). Thus, it seems plausible that participants may be generating their own unique appraisals about the events in the television show as they unfold. The results from our HMM analysis support this latter interpretation. We find that participants are more likely to occupy the same vmPFC state during the more emotionally evocative scenes and that distinct states appear to map onto valenced interpretations of the scene (e.g., good or bad), providing a direct replication of our fMRI study (Chang et al., 2021). Importantly, in these emotionally salient scenes, approximately only 50% of our sample appeared to occupy the same state. This likely explains why we did not observe synchronization using the ISC method, which would require more consistent temporal synchronization from the majority of the participants. Moreover, these results are also consistent with previous work that has found that the vmPFC may be involved in assessing the saliency of a particular event (Jafarpour et al., 2019).

More broadly, this work presents a fundamental challenge to studying the computations performed by the vmPFC. The majority of cognitive neuroscience research relies on group analyses to make inferences and these findings reveal that participants do not share consistent processes when viewing a rich and engaging naturalistic stimulus. Future work will need to develop novel research paradigms and analytic frameworks that can account for this intersubject heterogeneity to better characterize the computations performed by this region of cortex. Several promising approaches include the idiographic approach employed in studies of valuation (Rangel et al., 2008) and aesthetic experiences (Isik and Vessel, 2019), intersubject representational similarity analysis (Chen et al., 2020; Finn et al., 2020; van Baar et al., 2019), and fitting HMMs directly to single subject brain activity (Chang et al., 2021).

There are several potential limitations of this work that are important to highlight. First, it is unknown to what extent these results may be attributed to the pathophysiology of epilepsy or its pharmacological treatment. Both chronic refractory epilepsy and antiepileptic drugs could have a profound effect on the patient’s brain responses and cognitive functions (Brunbech and Sabers, 2002; Elger et al., 2004; Motamedi and Meador, 2003; Rantanen et al., 2011; van Rijckevorsel, 2006). However, we presume that these results will generalize beyond the specific participants and patient populations included in our analyses. Second, our analyses assume that our sampled electrodes in the vmPFC cover regions performing similar functions across participants. However, it is well known that the vmPFC is composed of several functionally separable regions (Kahnt et al., 2012). Thus, it is possible that the implanted electrodes may be located in regions that are not functionally homogeneous across participants. This particular issue may be more problematic for single unit recordings, as LFPs are likely reflecting coordinated firing of populations of neurons distributed across larger areas of cortex (Nir et al., 2007). Third, the maximum number of components we were able to estimate for the SRM was determined by the smallest number of electrodes implanted in any participant (Chen et al., 2015a). It is highly likely that the vmPFC contains many more than 6 latent components (the fewest number of electrodes in this region in a single participant). However, we think this is unlikely to change our results as we focused specifically on the component with the highest synchronization across participants. Fourth, our HMM analysis assumes that all participants experienced the same number of states. It is highly unlikely that all subjects experienced the same number of latent states, but this simplifying assumption was necessary to enable the alignment of states and compute the state concordance across participants.

In summary, using a naturalistic paradigm with sEEG recordings, we investigated intersubject synchronization of dynamic LFP signals in the vmPFC. Overall, we found minimal evidence indicating that signals in this region are shared across participants during this experimental context after removing artifacts related to spatial distance and within-subject and within-stereotactic strip clustering. However, our individual-HMM analysis revealed that some vmPFC states appeared to show increased alignment during emotionally salient scenes. These results are highly consistent with previous work using fMRI (Chang et al., 2021) and suggest that the vmPFC is involved in processing affective responses to ongoing experience in a state-like manner, but the specific states and temporal sequences are idiosyncratic to each individual. Beyond the vmPFC, we observed strong synchronization in the ***δ***,***α***, and ***β*** bands of auditory cortex that directly mapped onto properties of the eliciting auditory stimulus. In addition, we demonstrate that SRM (Chen et al., 2015a; Haxby et al., 2020) can be a promising technique to aid in functionally aligning signals from electrodes implanted in different regions across participants and also in denoising sEEG data.

## Methods

### Participants

In this study, we recruited 14 epilepsy patients (mean age = 38.6, sd=11.7; 5 females) undergoing inpatient stereo-EEG monitoring at the Dartmouth-Hitchcock Medical Center. sEEG electrode placement is determined solely based on clinical needs and we only included participants with electrodes implanted in both vmPFC and auditory regions (n=6, mean age = 45.6, sd = 12.3, 3 females), (See table 1 for detailed information on electrodes and subjects). All participants provided informed consent approved by the Institutional Review Board at Dartmouth-Hitchcock Medical Center.

### Experiment Procedure

Participants watched the first episode of the 45-minute television drama *Friday Night Lights* while undergoing stereo-EEG monitoring at their bedside. This show was selected because of its engaging and emotionally evocative content. The show was presented on a Windows computer using either PsychoPy (Peirce, 2007) or SuperLab (Cedrus, San Pedro, CA USA). Triggers indicating the onset and offset of the stimulus were sent to the sEEG recording system. Electrodes used in the study were either 0.86 mm diameter Ad-Tech (Ad-Tech Med Instr Corp, USA) sEEG with 10 independent recording contacts per electrode or 0.8 mm diameter PMT (PMT Corp., USA) sEEG with 8-12 independent recording contacts per electrode. Intracranial data were recorded on a Natus XLTek EMU 128 system at 2048Hz.

### Electrode Placement and Channel Selection

We obtained a pre-operative whole-brain T1-weighted MRI scan for each patient with a 3-T scanner using a 3D BRAVO sequence, at TR = 8.4 ms, TE = 3.4 ms, voxel size = 0.5 * 0.5 * 1.25 mm3. All scans were visually inspected for motion, abnormalities and other artifacts by certified radiologists at Dartmouth-Hitchcock Medical Center. Post-operative CT head scans were obtained for each patient at voxel size = 0.5 × 0.5 × 0.5 mm^3^ to compute the locations of each depth electrode. Both pre-operative MRI scans and post-operative CT scans were co-registered using SPM12 toolbox in Matlab 2017b. We determined the locations of depth electrodes by thresholding the registered CT image. Each depth electrode was represented by a cluster of dots in the registered CT image, while the coordinates were calculated by averaging all dots within each cluster. Individual T1-weighted MRI scans were normalized to Montreal Neurological Institute (MNI) space using the SPM 12 toolbox.

We assigned electrodes to each region of interest (ROI) based on broad areas of cortex functionally defined by the Neurosynth meta analytic database (Yarkoni et al., 2011). For each electrode we created a sphere of radius 2mm centered at the electrode’s MNI location. We then calculated the percentage overlap between the electrode sphere and each ROI region mask. We included electrodes with greater than 50% overlap within each ROI region mask. Electrode assignments and ROI region masks can be viewed at https://neurovault.org/collections/9709/).

### Signal Preprocessing

A careful visual inspection was conducted on the collected sEEG data by a qualified neurologist to exclude all bad/corrupted electrodes or electrodes with excessive epileptic activities. sEEG data was high-pass filtered at 0.1 Hz to remove slow drifts and a 60 Hz FIR notch filter was applied to remove electrical interferences. We used Laplacian re-referencing to electrodes on each stereotactic strip to extract local population-level activity (Li et al., 2018) and downsampled the data to 512 Hz to speed up subsequent analysis computations. Data preprocessing was performed in Python 3.7.9 with the MNE (Gramfort et al., 2013), Numpy (Harris et al., 2020), and Scipy (Virtanen et al., 2020) packages.

### Time-Frequency Processing

Unless otherwise specified, we used Morlet Wavelet Transforms implemented in the MNE package (function: *tfr_array_morlet) (Gramfort et al*., *2013)* with a width of 7 cycles for all time frequency decomposition analyses discussed in the paper. Specifically, we created 30 logarithmically spaced frequency bins ranging from 1 Hz to 150 Hz, applied the wavelets to each frequency bin, and calculated the power. Power modulations that fall between 56 - 64 Hz and 116-124 Hz are not included because of their proximity to the 60 Hz power line noises and 120 Hz harmonics. We divided the power in each frequency bin by its mean to convert the absolute power values to a proportion of the mean (Ossandón et al., 2011). This normalization procedure minimizes power differences across frequency bands resulting from their 1/f distribution. Finally we averaged all power estimates for the wavelet frequency bins within (1-2Hz), ***θ*** (2-5 Hz), ***α*** (5-14 Hz), ***β*** (14-30 Hz), low ***γ*** (30-70 Hz), high ***γ*** (70-150 Hz), or overall broadband power (1-150Hz) bands. We also ran all time-frequency decomposition analyses with Hilbert transformation and observed similar results.

### Power-Spectrum Density

We calculated the power spectrum density (PSD) of the signal using Welch’s Method with a Hamming window of 1s with 50% overlap. In Fig 4B & Fig S3, we removed the 1/f power decay by log-transforming the power and frequency bins estimated from Welch’s Method, fitting a linear regression and finally plotting the residuals from the regression. We used implementations from mne-python (function: *psd_array_welch*) (Gramfort et al., 2013) for Welch’s Method and implemented the linear regression with scikit-learn (function: *LinearRegression*) (Pedregosa et al., 2011)

### Inter-subject Correlation (ISC)

In order to measure the degree of shared information patterns across subjects, we used the *Intersubject correlation* (ISC) metric (Hasson et al., 2004). ISC is a simple yet effective way to reliably extract the shared brain responses to complex naturalistic stimuli across individuals (Nastase et al., 2019). ISC was originally developed for fMRI data analysis but has also been applied to other neuroimaging modalities such as EEG (Dmochowski et al., 2012; Imhof et al., 2020; Kaneshiro et al., 2020; Maffei, 2020; Poulsen et al., 2017), MEG (Chang et al., 2015; Chen and Farivar, 2020; Thiede et al., 2020).

We applied ISC to our intracranial EEG data to study the degree of functional alignment across subjects separately within auditory and vmPFC electrodes. We calculated ISC as the mean value of the lower triangle of the pairwise correlation matrix, excluding the diagonal elements. We applied fisher’s z-transform to transform the pairwise correlation value prior to taking the mean and used the inverse fisher transform to convert the mean z-transformed value back to correlation. Each correlation pair is calculated as:

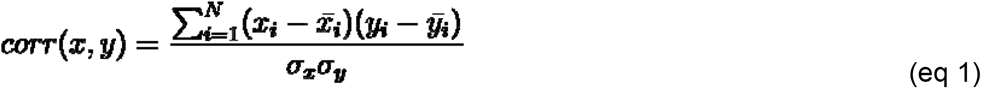

where,

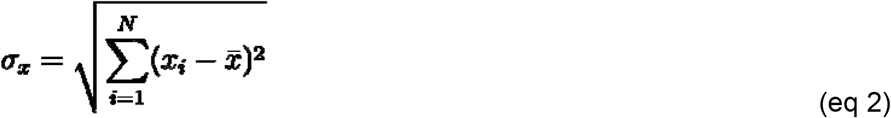

and

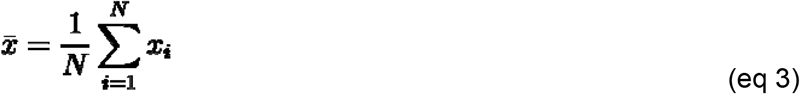

The overall ISC is thus,

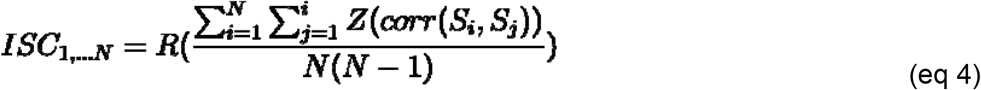

where,

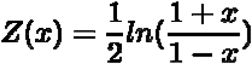

and

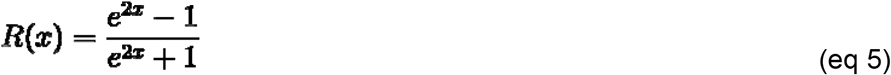

***S*** is the ‘subject’ matrix of shape either number of electrodes by time (for raw electrode & residualized analysis, N=44) or number of subjects by time (for the subsequent SRM analysis, N=6).

To test if ISC values are significantly above zero, we used subject-wise bootstrapping as recommended by (Chen et al., 2016). Specifically, we sample the pairwise correlations in the lower triangle correlation matrix with replacement and calculate the ISC after removing correlations between the same electrode. We repeated the process 5000 times and calculated the p-value for our two-tailed hypothesis test. We used the ISC and bootstrap implementations from our python analysis package nltools (Chang et al., 2018) available at https://nltools.org/.

### Distance Regression

To perform our distance regression (Fig. 2A), we first computed the pairwise correlation matrix separately for vmPFC or auditory electrodes across 6 subjects. Next, we calculated a pairwise Euclidean Distance matrix using the x/y/z millimeter MNI coordinates separately for vmPFC or auditory electrodes across 6 subjects. Prior to running the regression, we Fisher z-transformed the correlation matrix and centered the distance matrix. We then added within-subject dummy matrices for each subject, in which electrode pairs clustered within a subject were indicated by a value of 1, and 0 everywhere else. In addition, we added within-stereotactic strip dummy matrices, to indicate whether electrodes were located on the same strip within a subject with a 1 or 0. We estimated a separate linear model for each region of interest (e.g., vmPFC or auditory cortex) using the pairwise correlation matrix as the response variable and the euclidean distance matrix, 6 dummy subject matrices and dummy electrode matrix (10 for auditory while 16 for vmPFC) as explanatory variables using the function ‘lm’ in R (Computing and Others, 2013). We did not include a global intercept for the regression. The residuals of the model are obtained and used in subsequent ISC analyses. We calculated the variance explained using a nested model comparison approach where we compared the total variance explained to a subset of the model omitting either the spatial distance or the within-subject clustering dummy indicator matrices.

### Shared Response Model

We performed functional alignment (Haxby et al., 2020) using the Shared Response Model (SRM; (Chen et al., 2015a)) separately on auditory and vmPFC electrodes across 6 subjects. As demonstrated in Fig. 4, SRM is a matrix factorization model that decomposes the electrode-by-time matrix for each subject into a common shared component matrix and an orthogonal subject-specific basis matrix. The objective function is to minimize the reconstruction loss across all 6 subjects. SRM identifies common activity patterns that are present across subjects and provides a method to transform the original electrode activities into a shared latent component space. The number of features ***k*** in our study is chosen as the minimum number of electrodes present in any given participant (k=5 for auditory, k=6 for vmPFC).

For each subject’s electrodes activity matrix ***X***_***i***_, i=1..6, of shape **(*n***_***i***_,***t*)** where ***n***_***i***_ is the number of electrodes present in the subject (see Table S1) and ***t*** is the time length of the sEEG recording (∼45 minutes), we decompose ***X***_***i***_ **≈ *W***_***i***_***S***, where ***W***_***i***_ is the orthogonal subject-specific basis function of shape **(*n***_***i***_,***k*)** satisfying 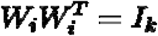 and ***S*** is the shared components matrix of shape ***(k***,***t)***. The objective function is thus to minimize the squared reconstruction loss as 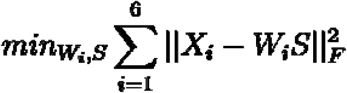, where **‖ · ‖**_***F***_ is the frobenius norm. (see details in (Chen et al., 2015b) about how to computationally derive a solution to the objective function). After we have identified the ***W***_***i***_ for each subject and ***S*** for all subjects, we can use the subject’s reconstructed activity matrix 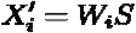, of shape **(*k***,***t*)**, to approximate the subject’s raw electrode activity matrix ***X***_***i***._ For each component ***m*=1**..***k***, we selected the ***m − th*** row (which leads to a matrix of shape **(1**,***t*)** for each subject) of the reconstructed matrix 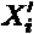 across 6 subjects and calculated the ISC. We selected the component ***m*** that yields the maximum ISC across 6 subjects. We used a modified version of the original Brainiak version of SRM implemented in our nltools package (Chang et al., 2018).

### Statistical Tests

All hypothesis tests comparing regions used non-parametric sign permutation tests. In order to test the significance of differences between auditory and vmPFC ISC values, we randomly shuffled the group assignment of each pairwise correlation and calculated the ISC differences for the re-shuffled pairwise correlations. We repeated the process 5000 times to generate a null distribution and calculated p-values for our two-tailed hypothesis test, by counting the number of iterations that our observed difference exceeded the null distribution. For one-sample tests, we randomly multiplied each value by [1,-1] and generated a null distribution using 5000 iterations. To test if ISC values are significantly above zero, we used subject-wise bootstrapping as recommended by (Chen et al., 2016) and described above in ISC section.

### Extracting audio power envelope

We extracted the power envelope of the auditory stimulus using a similar procedure as (Honey et al., 2012). We extracted power from the auditory channel using multi-tapering where power is estimated every 200 Hz from 200 Hz to 5000 Hz (25 frequency bands in total) with 3 tapers. We took the logarithm of each frequency band and further z-score normalized the power in each frequency band. Finally we averaged the powers across 25 frequency bands to get the final power envelope for the audio stimuli.

### Hidden Markov Model

Hidden Markov Models are generative probabilistic state-space models which assume that the current sequence of observations is generated by a sequence of discrete latent state variables. The transitions between latent states are assumed to follow the first order Markov Process, where the probability of transition to the next immediate state depends only on the most recent state. We have also made the implicit assumption that the observed variables follow a multivariate Gaussian distribution with diagonal covariance matrix.

We fit an individual HMM separately on each participant’s broadband power SRM components. Specifically we first applied a 6-component SRM to the raw vmPFC electrodes across 6 subjects, and obtained the resulting transformed matrix of shape **(6**,***t*)** for each subject. We extracted the broadband power from the SRM components for each participant and normalized the power for each SRM component for each participant. We fit an HMM to the normalized broadband power matrix for each subject using the *hmmlearn Package* (version 0.2.5). We used the Viterbi algorithm to maximize a posteriori probability estimates and infer the most likely sequence of latent states. Model parameters were estimated with expectation-maximization. In order to determine the number of states *k* for our HMM model, we calculated the Bayesian Information Criterion (BIC) value over a range of ***k* = [2**,**15]**. BIC finds the simplest model to fit the data by balancing the maximum likelihood function and the model parameters & number of data points.

Specifically BIC is calculated as:

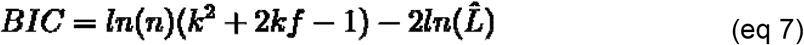

where ***n*** is the number of observations,***k*** is the number of parameters in the model, ***f*** is the number of features (which equals to 6 SRM components in our case), and is 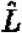 the maximum log-likelihood of the HMM model fit. We calculated the BIC value for each chosen ***k* = [2**,**14]** for each participant. For each state we averaged the BIC across 6 participants. We then selected the ***k*** which gives the maximum reduction in BIC value (i.e., the derivative of the BIC with respect to ***k***; Fig. S7). For each fitted model, we aligned the latent states across participants by maximizing the correlation similarity between the model pairs using the Hungarian Algorithm (Kuhn, 2005).

### Code

All code used to perform these analyses are available on Github https://github.com/cosanlab/naturalistic_iEEG_func_align/.

## Acknowledgments

This work was supported by awards from the National Institute of Mental Health R01MH116026 and R56MH080716 to LJC. TKX is supported by the Big Data in the Life Sciences Training Program by Burroughs-Wellcome Fund.

