## Supplemental Files for "Minimal functional alignment of ventromedial prefrontal cortex intracranial EEG signals during naturalistic viewing"

#### Time-Frequency Intersubject Correlation Analyses

In addition to the ISC analyses performed on broadband power, we also computed ISC separately for each canonical frequency band separately for the auditory and vmPFC regions of interest. First, we computed the time-frequency ISC analysis on each frequency band from the filtered electrode time series using the ISC and wavelet frequency decomposition approaches described in the methods section. Overall, we found that both auditory and vmPFC ISC values were statistically significantly different from zero, but not from each other (Fig. S1). Second, we performed the same time-frequency ISC analyses using the residualized data from our distance regression procedure and found that none of the vmPFC frequency bands significantly differed from zero, but the auditory cortex exhibited significantly greater ISC in the
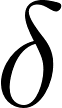
<https://www.codecogs.com/eqnedit.php?latex=%5Cdelta#0>,
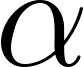
<https://www.codecogs.com/eqnedit.php?latex=%5Calpha#0>,
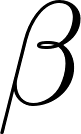
<https://www.codecogs.com/eqnedit.php?latex=%5Cbeta#0>,
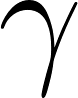
<https://www.codecogs.com/eqnedit.php?latex=%5Cgamma#0>, and broadband power (Fig. S2). Third, we performed the time-frequency ISC analysis on the SRM component that exhibited the highest synchrony across participants for the auditory and vmPFC cortex. The results are described in the main text (Fig. 5).

#### Denoising sEEG data with the Shared Response Model (SRM)

The SRM attempts to identify latent components that are shared across participants. An underappreciated aspect of this procedure is that it will remove signals that are not shared across participants. In intracranial data, there are many sources of noise that are unique to each participant and can be filtered out using this approach. We outline several supplementary analyses we performed to evaluate the SRM’s ability to remove different types of known artifacts present in the data such as the 60Hz line noise, interictal epileptiform discharges (IEDs) and Ictal (seizure) activities.

##### Power Line Noise

The 50Hz or 60Hz power line interference is one of the most prevalent sources of noise in EEG/MEG studies (Leske and Dalal, 2019). Although power line interferences are concentrated in the 50 or 60 Hz frequency bands, they are not typically phase locked across participants. Because we fit the SRM to data in the time-domain, we anticipate that this procedure may mitigate the 60Hz power line interference. To test this hypothesis, we conducted SRM on the non-preprocessed raw electrode data for each subject and investigated the power spectrum density of the resulting SRM components. Overall, we found that SRM was able to reduce the line noise signal, but was unable to completely remove it in most participants. We quantitatively tested the effect of power line noise removal for each subject by first calculating the proportion of power in bands 58-62 Hz & 118-122 Hz separately for each raw unfiltered auditory electrode
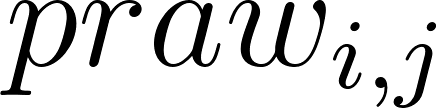
<https://www.codecogs.com/eqnedit.php?latex=praw_%7Bi%2Cj%7D#0> and the SRM components
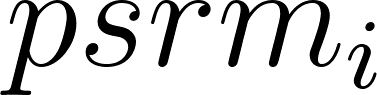
<https://www.codecogs.com/eqnedit.php?latex=psrm_%7Bi%7D#0>, where
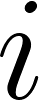
<https://www.codecogs.com/eqnedit.php?latex=i#0> indicates the subject number and
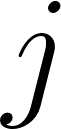
<https://www.codecogs.com/eqnedit.php?latex=j#0> indicates the participant’s electrode id. We then averaged the proportion for all raw electrodes within a subject
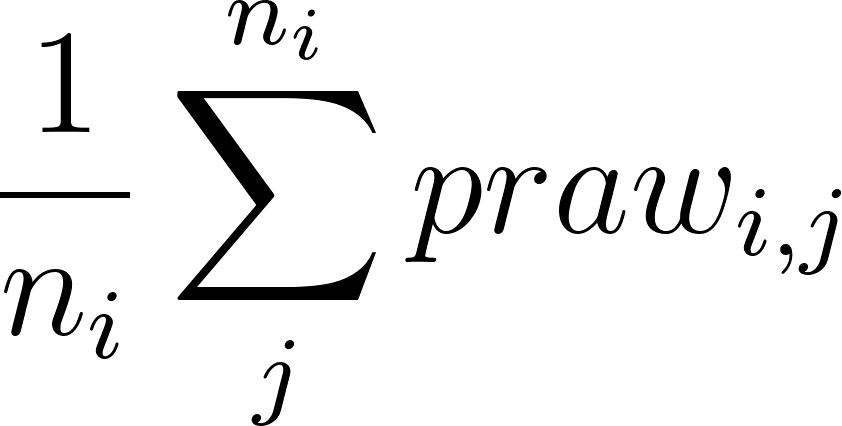
<https://www.codecogs.com/eqnedit.php?latex=%5Cfrac%7B1%7D%7Bn_i%7D%20%5Csum_j%5E%7Bn_i%7D%20praw_%7Bi%2Cj%7D#0>. We repeated the procedure to obtain the averaged power line noise reduction across 6 subjects and find that the SRM procedure is able to reduce power line noise by an average of 51% across 6 subjects, but this only approached statistical significance using a non-parametric paired permutation test (p=0.0625). The non-parametric paired permutation test is performed by first calculating the ground averaged difference between SRM and raw proportions for all 6 (subject) pairs. We then permute every possible pair (in total 2^6 = 64 permutations) by swapping SRM and raw proportions and re-calculated the averaged difference between the permuted SRM and raw proportions for all 6 pairs. The exact p-value is the number of times the permuted averaged difference exceeds the observed averaged difference divided by the total number of all possible permutations (64).

##### Interictal Epileptiform Discharges and Ictal Activity

Interictal epileptic discharges (IEDs) and Ictal (seizure) activities are common sources of noise in intracranial EEG data. IEDs are spikes or spike-waves that occurred independently from seizures and are thought to indicate pathological alterations from the normal neuronal activities (Chatrian et al., 1974; GOTMAN and J, 1980; Keller et al., 2010; Kooi, 1966; Walczac and Jayakar, 1997). IEDs and Ictal activities do not occur in time across subjects, thus we hypothesized that SRM should be able to remove much of these artifacts. We tested this hypothesis in two different ways. First, we performed a qualitative analysis, in which we selected 20 second time intervals when the electrodes exhibit visually salient IED with ictal patterns. We selected 3 electrodes (all from Left Amygdala) for each of the 5 subjects (s1-s5, s6 does not exhibit salient IED or ictal activities across all channels). Note that the start time and end time of the 20 second time interval is fixed for the 3 electrodes within the same subject but not across subjects (because it is not possible for all subjects to simultaneously have IED and Ictal activities). We then applied 3-component SRM to the electrodes across 5 subjects and selected the component that maximizes intersubject synchrony. As demonstrated in Fig. 4C the SRM successfully removed the IED and Ictal activities.

Second, we performed a more controlled simulation experiment to quantify the degree to which SRM can remove noise that is idiosyncratic to a specific participant. Simulations can allow us to quantify how much of the artifact SRM can remove when we know the ground truth. This is particularly important as it is currently difficult to reliably identify IEDs or Ictal activities using automated methods (Keller et al., 2010; Thomas et al., 2018). In this simulation, we iteratively added different levels of Gaussian noise to the original auditory electrodes to examine the ability of SRM in extracting meaningful signals in presence of noise. For each subject we simulate independent Gaussian noise for each time point using 3 different signal-to-noise ratios (SNR): 0.01 (low), 0.25 (medium) & 1 (high) and add this noise to all electrodes to create simulated contaminated electrodes within each subject. We then applied SRM on all contaminated electrodes across 6 subjects and obtained the maximally aligned SRM component. In order to quantitatively measure to what extend SRM can remove the simulated noise, for each subject we correlated the noise with every contaminated electrode
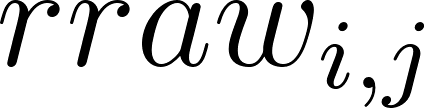
<https://www.codecogs.com/eqnedit.php?latex=rraw_%7Bi%2Cj%7D#0>,where
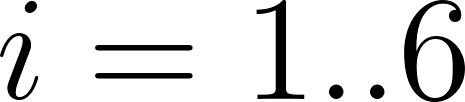
<https://www.codecogs.com/eqnedit.php?latex=i%3D1..6#0> represents subject and j = 1..
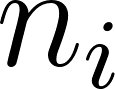
<https://www.codecogs.com/eqnedit.php?latex=n_i#0> represents the electrode within each subject, and the SRM component
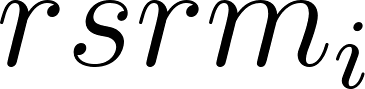
<https://www.codecogs.com/eqnedit.php?latex=rsrm_i#0>. We averaged the noise correlations for all raw electrodes within a subject
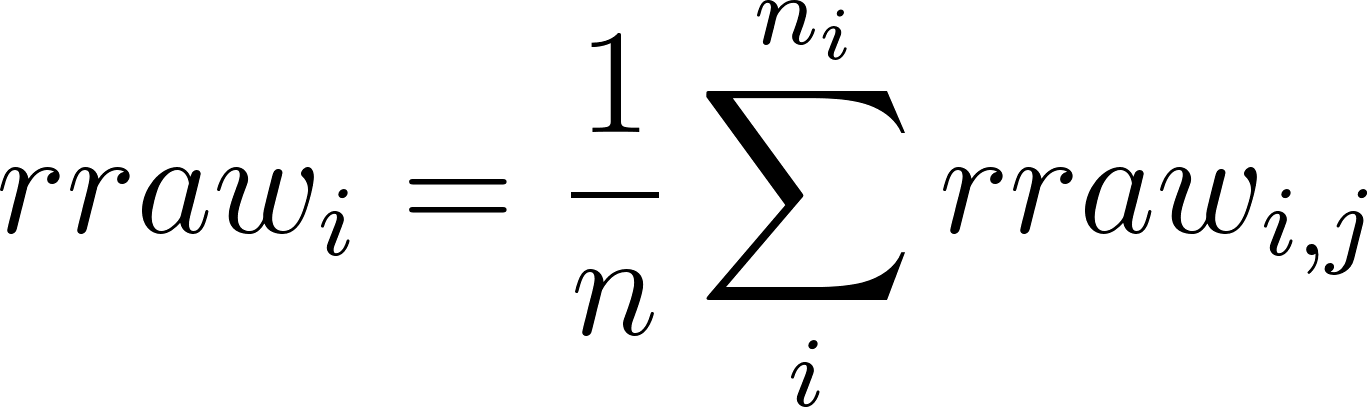
<https://www.codecogs.com/eqnedit.php?latex=rraw_%7Bi%7D%20%3D%20%5Cfrac%7B1%7D%7Bn%7D%5Csum_i%5E%7Bn_i%7D%20rraw_%7Bi%2Cj%7D#0>We then calculated the proportion of noise removed by SRM for each subject as
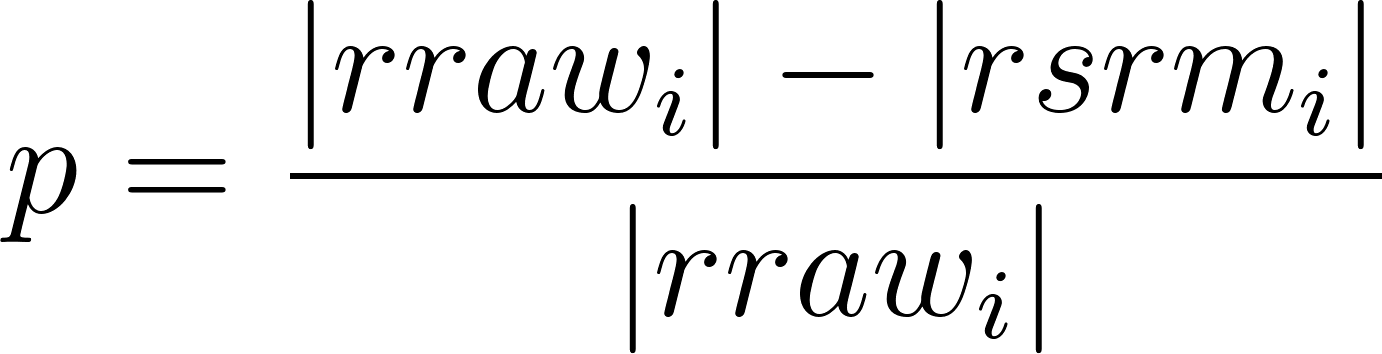
<https://www.codecogs.com/eqnedit.php?latex=p%20%3D%5Cfrac%7B%7Crraw_i%7C-%7Crsrm_i%7C%7D%7B%7Crraw_i%7C%7D#0>. We ran 2,500 simulations for each SNR value and found that across every SNR value SRM can significantly reduce the noise by an average of 77.9% (p < 0.001 for all SNRs, see Fig. S5). This indicates that SRM was able to significantly reduce the artifactual signal. To ensure that the signal was no longer contaminated by the artifactual signal, we decomposed the max SRM component from the data with the simulated noise to the
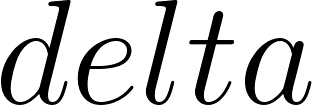
<https://www.codecogs.com/eqnedit.php?latex=delta#0> band (1-2 Hz) with Morlet Wavelets and correlated it with the audio and found that the SRM component still strongly corresponds to the audio stimuli across all SNR levels, (SNR 0.01: r=-0.18, p<0.001),(SNR 0.25: r=-0.19, p<0.001), (SRN 1: r=-0.20, p<0.001). Note that we focused on the
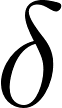
<https://www.codecogs.com/eqnedit.php?latex=%5Cdelta#0> band because we have already observed strong negative coupling effects as illustrated in Fig. 4 A.

### Supplementary Tables

**Table S1.** Number of electrodes implanted in each region of interest for each included participant.

| Subject ID | # Auditory Electrodes | # vmPFC electrodes |
| --- | --- | --- |
| s1 | 5 | 11 |
| s2 | 8 | 10 |
| s3 | 8 | 12 |
| s4 | 5 | 6 |
| s5 | 5 | 11 |
| s6 | 13 | 6 |
| Totals | 44 | 56 |

### **Table S2**. Fitted residual regression model for auditory electrodes

| **Variable** | **Regressor** | **Estimate** | **Standard Error** | **T** | **P - Value** |
| --- | --- | --- | --- | --- | --- |
| Spatial Distance |  |  |  |  |  |
|  | Euclidean Distance | 0.00 | 0.00 | -2.65 | 0.008 |
| Subject Indicator |  |  |  |  |  |
|  | Subject 0 | 0.92 | 0.05 | 19.81 | < 0.001 |
|  | Subject 1 | 1.14 | 0.03 | 42.49 | < 0.001 |
|  | Subject 2 | 0.55 | 0.02 | 23.61 | < 0.001 |
|  | Subject 3 | 1.67 | 0.03 | 56.45 | < 0.001 |
|  | Subject 4 | 0.25 | 0.05 | 5.35 | < 0.001 |
|  | Subject 5 | 0.47 | 0.01 | 37.57 | < 0.001 |
| Strip Indicator |  |  |  |  |  |
|  | 1RAHCD | 0.48 | 0.06 | 8.07 | < 0.001 |
|  | 2RPTD | 0.42 | 0.04 | 11.80 | < 0.001 |
|  | 2LPTD | 0.67 | 0.10 | 6.91 | < 0.001 |
|  | 3RSTGD | 0.42 | 0.05 | 9.37 | < 0.001 |
|  | 3RPTD | 0.63 | 0.05 | 14.14 | < 0.001 |
|  | 5RAHD | 0.96 | 0.06 | 16.06 | < 0.001 |
|  | 6LWD | 0.33 | 0.03 | 10.41 | < 0.001 |
|  | 6LMSTGD | 0.49 | 0.06 | 8.88 | < 0.001 |
|  | 6LPSTGD | 0.15 | 0.03 | 4.60 | < 0.001 |

**Table S3**. Fitted residual regression model for vmPFC electrodes

| **Variable** | **Regressor** | **Estimate** | **Standard Error** | **T** | **P - Value** |
| --- | --- | --- | --- | --- | --- |
| Spatial Distance |  |  |  |  |  |
|  | Euclidean Distance | 0.00 | 0.00 | -2.30 | 0.022 |
| Subject Indicator |  |  |  |  |  |
|  | Subject 0 | 0.89 | 0.01 | 73.81 | <0.001 |
|  | Subject 1 | 1.39 | 0.01 | 100.76 | <0.001 |
|  | Subject 2 | 0.72 | 0.01 | 62.30 | <0.001 |
|  | Subject 3 | 1.49 | 0.03 | 43.25 | <0.001 |
|  | Subject 4 | 0.19 | 0.01 | 16.36 | <0.001 |
|  | Subject 5 | 0.58 | 0.03 | 21.18 | <0.001 |
| Strip Indicator |  |  |  |  |  |
|  | 1LOFCD | 0.33 | 0.03 | 12.02 | <0.001 |
|  | 1ROFCD | 0.43 | 0.05 | 9.36 | <0.001 |
|  | 1RACD | 0.85 | 0.08 | 10.88 | <0.001 |
|  | 2ROFD | 0.82 | 0.05 | 17.61 | <0.001 |
|  | 2RACD | 0.17 | 0.08 | 2.18 | 0.0294 |
|  | 2LOFD | 0.64 | 0.03 | 22.65 | <0.001 |
|  | 3LOFCD | 0.12 | 0.02 | 5.11 | <0.001 |
|  | 3ROFCD | 0.27 | 0.03 | 8.09 | <0.001 |
|  | 3RACD | 0.61 | 0.08 | 7.89 | <0.001 |
|  | 4LOFD | 0.00 | 0.04 | 0.01 | 0.992 |
|  | 5LOFCD | 0.26 | 0.03 | 7.82 | <0.001 |
|  | 5ROFCD | 0.87 | 0.08 | 11.20 | <0.001 |
|  | 5LACD | 0.09 | 0.05 | 1.98 | 0.0485 |
|  | 5RACD | 0.52 | 0.08 | 6.62 | <0.001 |
|  | 6LOFCD | 0.22 | 0.04 | 5.24 | <0.001 |
|  | 6LCD | 0.59 | 0.08 | 7.26 | <0.001 |

### Supplementary Figures

**Figure S1. Raw electrode Intersubject Correlation (ISC) by vmPFC/auditory by frequency bands.** In this analysis, we perform ISC on raw electrodes across 6 subjects after extracting power from different frequency bands. Error bars indicate 95% confidence intervals for the ISC values (See details in Method section) and the significance labels indicate whether there are significant differences in the ISC values between auditory and vmPFC electrodes within the same frequency bin. Overall, we found that ISC values are significantly higher than zero for both vmPFC and auditory electrodes across all frequency bands. However, there is no significant difference between vmPFC electrode ISC and auditory electrode ISC across all frequency bands.

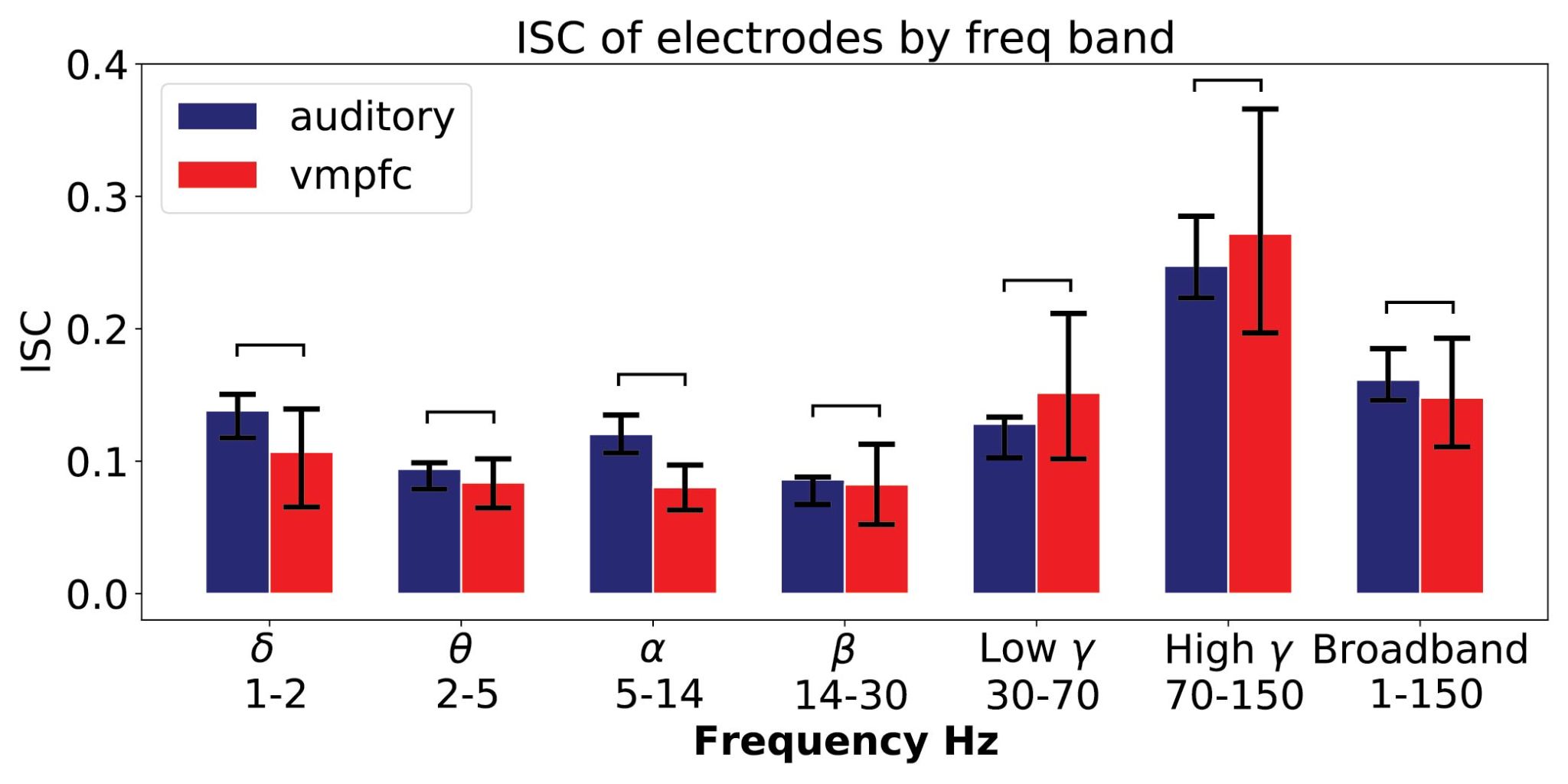

**Figure S2. Residualized ISC by vmPFC/auditory by frequency bands.** In this analysis, we examine differences in ISC between auditory cortex and vmPFC after removing variance associated with spatial distance and intersubject clustering. Error bars indicate 95% confidence intervals for the ISC values (see details in Method section) and the significance * labels indicate whether there are significant differences in the ISC values between auditory and vmPFC electrodes within the same frequency bin (*: 0.01<=p<0.05; **: 0.001<=p<0.01; *** : p<0.001). Overall, we observe poor cross subject alignment in vmPFC across multiple frequency bands. In contrast, we observe strong cross subject alignment in auditory across multiple frequency bands (significant: low
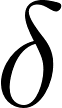
<https://www.codecogs.com/eqnedit.php?latex=%5Cdelta#0>,
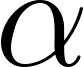
<https://www.codecogs.com/eqnedit.php?latex=%5Calpha#0>,
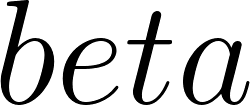
<https://www.codecogs.com/eqnedit.php?latex=beta#0> & Broadband power). The differences between electrode and vmPFC are significant in low
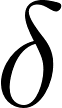
<https://www.codecogs.com/eqnedit.php?latex=%5Cdelta#0>,
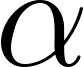
<https://www.codecogs.com/eqnedit.php?latex=%5Calpha#0>,
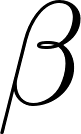
<https://www.codecogs.com/eqnedit.php?latex=%5Cbeta#0>,
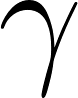
<https://www.codecogs.com/eqnedit.php?latex=%5Cgamma#0> and broadband powers.

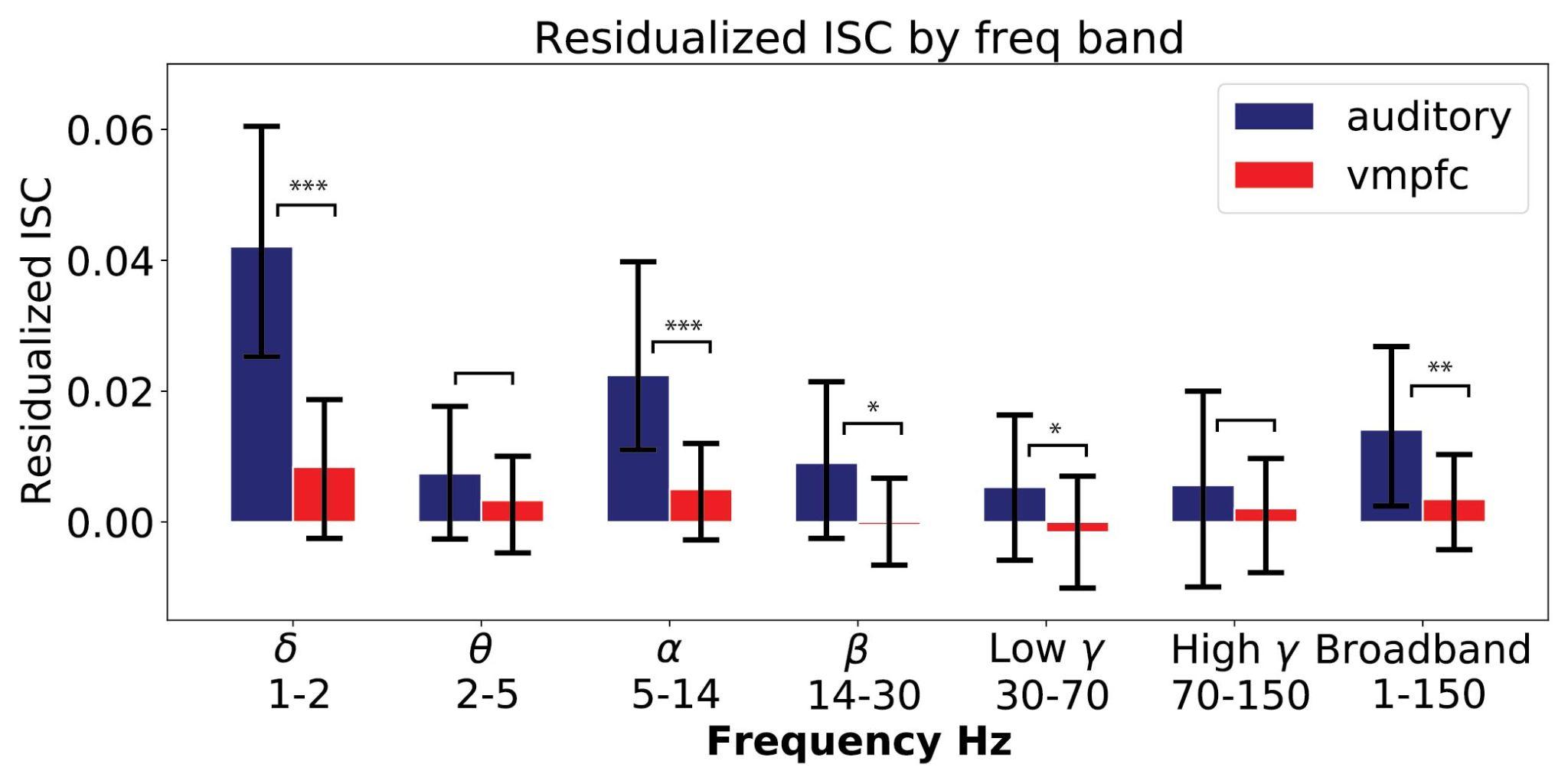

**Figure S3. Single subject power spectral density (PSD).** To estimate the extent to which SRM can reduce power line noise, we apply SRM to non-preprocessed raw sEEG auditory electrodes and plot the power spectrum density for each subject. We found that SRM is capable of reducing power line noises, but not completely removing them.

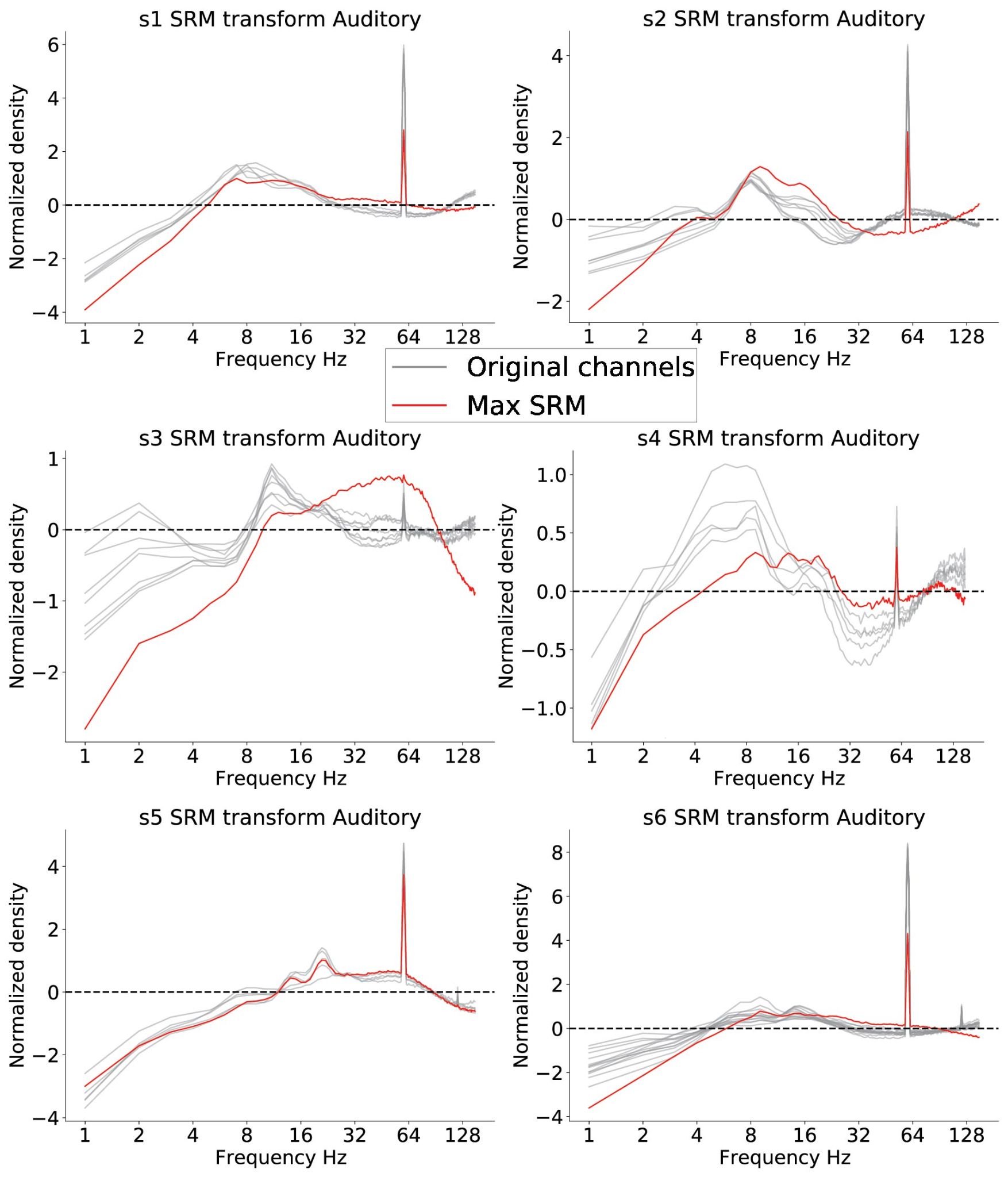

**Figure S4. Cross-validated SRM.** To rule out any bias in ISC values resulting from fitting and evaluating the SRM model on the same data, we performed a split-half cross-validation procedure. We plot broadband power (1-151 Hz) and power extracted from narrow frequency bands on the test data that is independent of the SRM training procedure. We see no appreciable differences in training or testing ISC for both auditory and vmPFC SRM components.

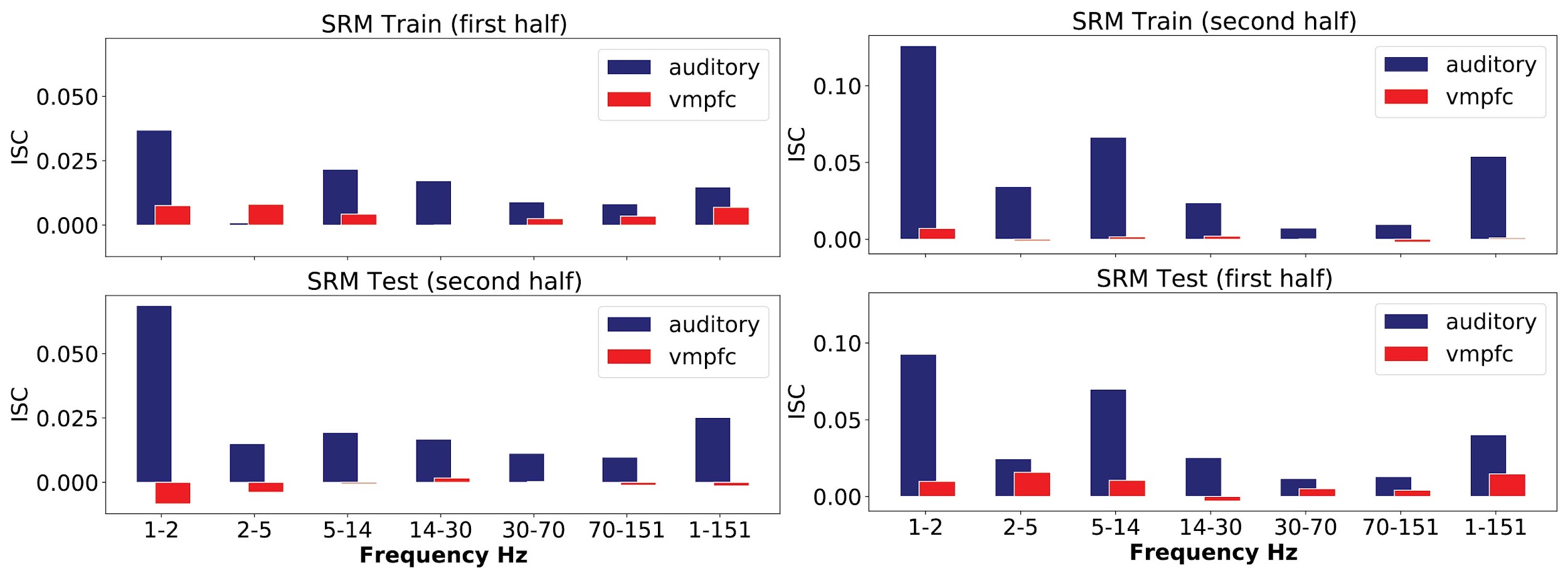

**Figure S5. SRM denoising effectiveness.** We simulated Gaussian noise at 3 different SNR ratios and found that SRM can significantly reduce the simulated noise across multiple SNRs. Error bars indicate 95% confidence intervals for the simulated noise reductions in SRM.

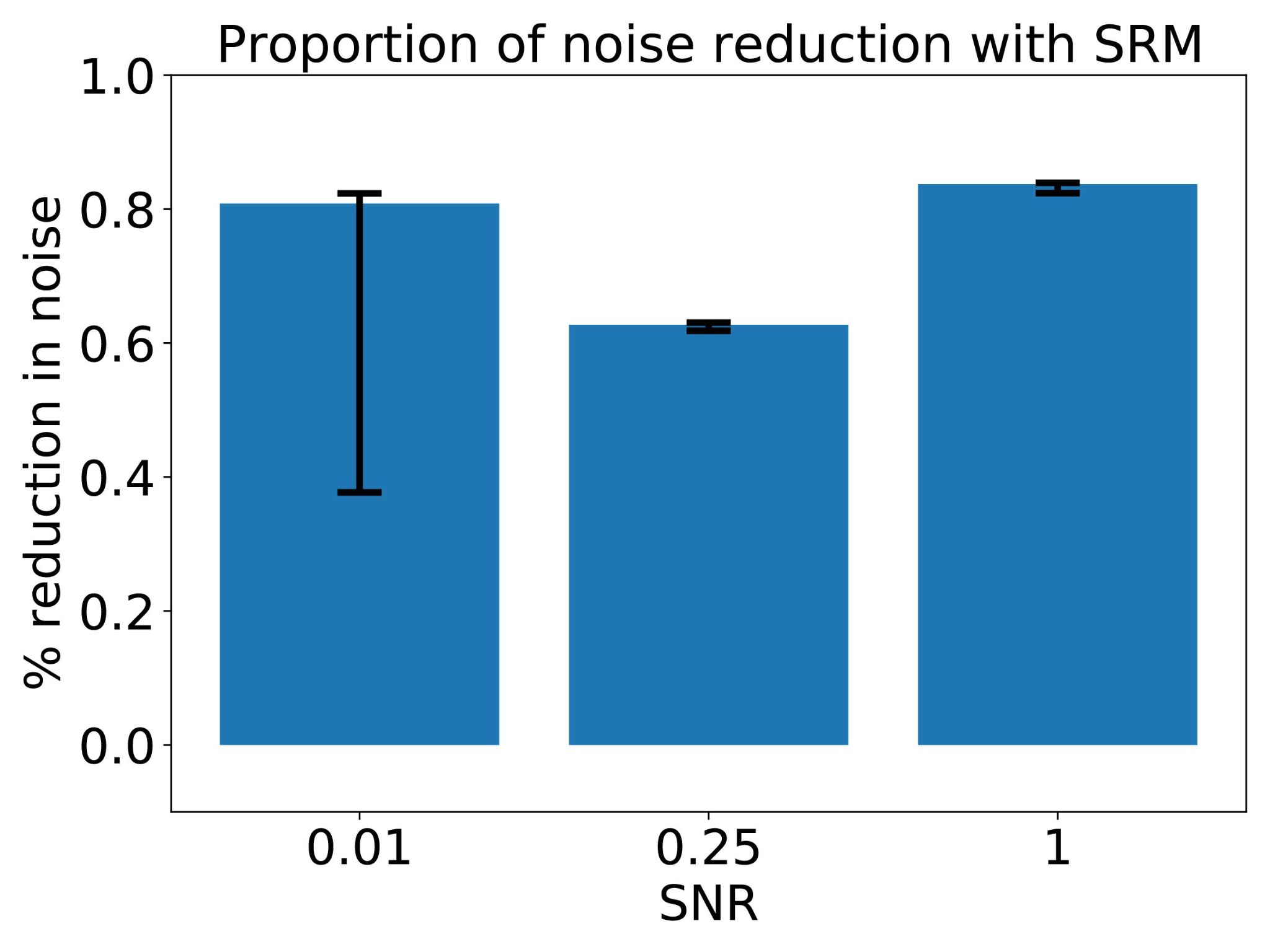

**Figure S7. HMM model fits.** Here we plot the subject-averaged Bayesian Information Criterion (BIC) for
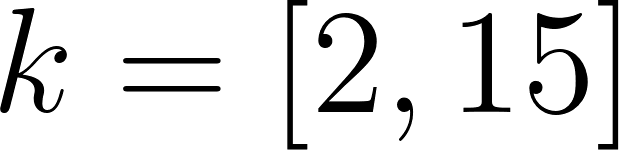
<https://www.codecogs.com/eqnedit.php?latex=k%20%3D%20%5B2%2C15%5D#0>. We fit a separate HMM model for each subject and averaged the BIC values across subjects for each
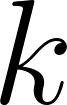
<https://www.codecogs.com/eqnedit.php?latex=k#0>. In panel A the bold solid blue line shows the averaged BIC by state number
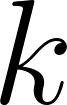
<https://www.codecogs.com/eqnedit.php?latex=k#0> while the other blue lines indicate individual HMM BIC. In panel B the bold solid line shows the first derivative of the averaged BIC by state number
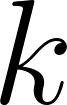
<https://www.codecogs.com/eqnedit.php?latex=k#0> while the other blue lines indicate individual HMM BIC derivative. The red dotted line indicates the number of state (k=3) that resulted in the largest drop in BIC.

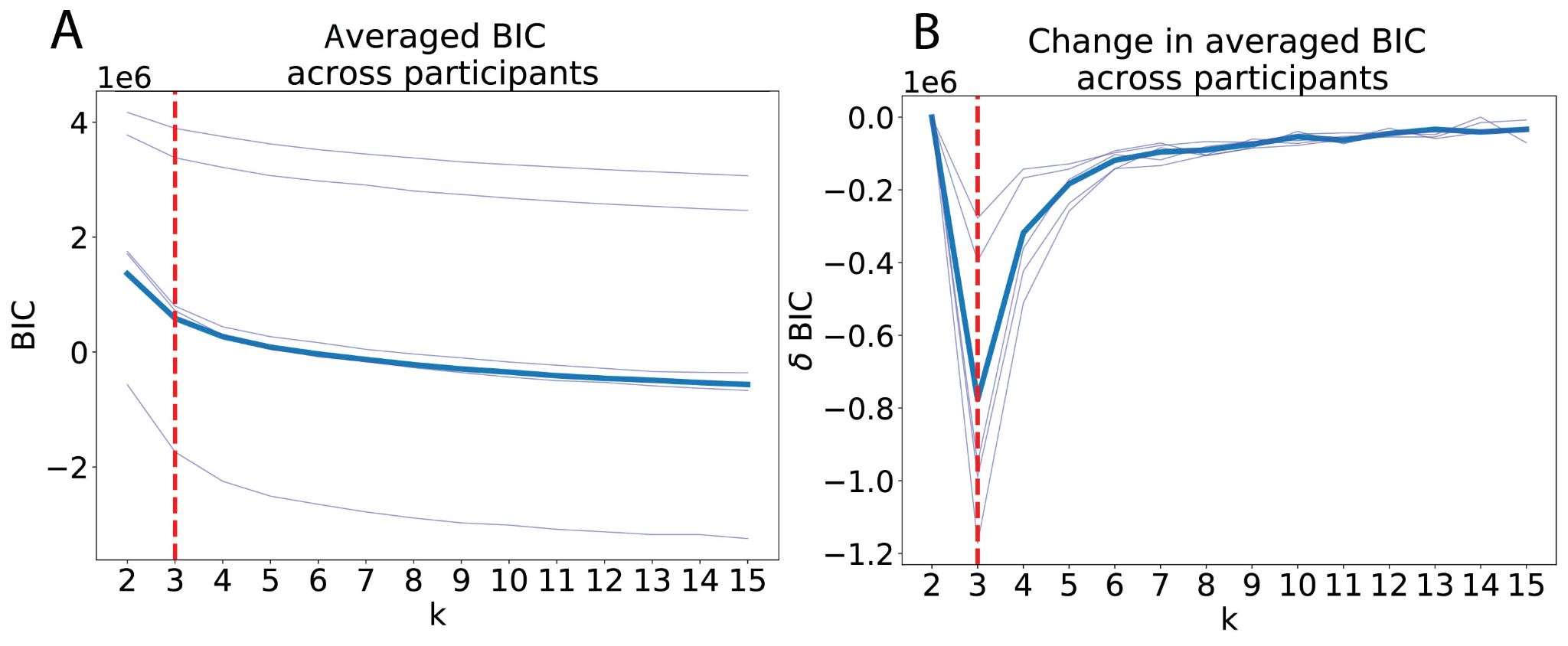

**Figure S8. Individual SRM PSDs.** Here we plot the log-log normalized power spectrum density for each subject’s max SRM component per auditory (left) or vmPFC (right) group. The 1/f trend is removed by fitting a first-order linear regression and obtaining the residuals.

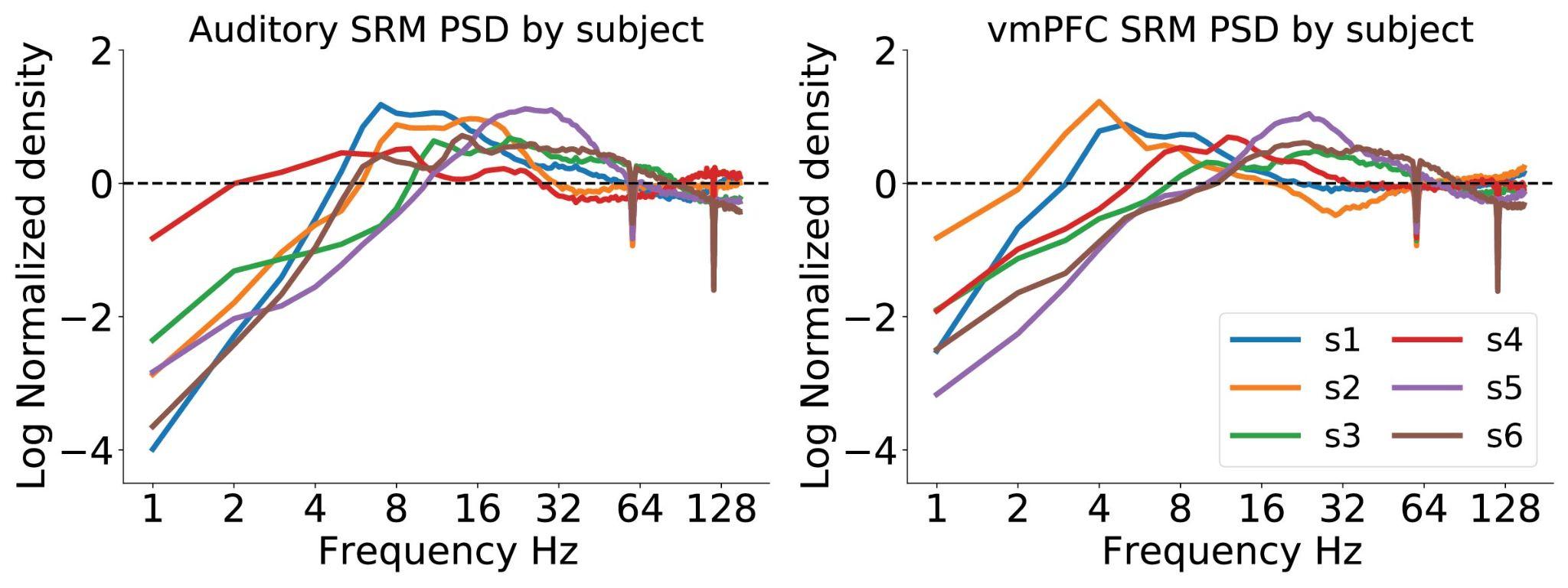
